# Homologs of circadian clock proteins impact the metabolic switch between light and dark growth in the cyanobacterium *Synechocystis* sp. PCC 6803

**DOI:** 10.1101/2021.04.21.440780

**Authors:** Nina M. Scheurer, Yogeswari Rajarathinam, Stefan Timm, Christin Köbler, Joachim Kopka, Martin Hagemann, Annegret Wilde

## Abstract

The putative circadian clock system of the facultative heterotrophic cyanobacterial strain *Synechocystis* sp. PCC 6803 comprises the following three Kai-based systems: a KaiABC-based potential oscillator that is linked to the SasA-RpaA two-component output pathway and two additional KaiBC systems without a cognate KaiA component. Mutants lacking the genes encoding the KaiAB1C1 components or the response regulator RpaA show reduced growth in light/dark cycles and do not show heterotrophic growth in the dark. In the present study, the effect of these mutations on central metabolism was analyzed by targeted and nontargeted metabolite profiling. The strongest metabolic changes were observed in the dark in Δ*rpaA* and, to a lesser extent, in the Δ*kaiAB1C1* mutant. These observations included the overaccumulation of 2-phosphoglycolate, which correlated with the overaccumulation of the RbcL subunit in the mutants, and taken together, these data suggest enhanced RubisCO activity in the dark. The imbalanced carbon metabolism in the Δ*rpaA* mutant extended to the pyruvate family of amino acids, which showed increased accumulation in the dark. Hence, the deletion of the response regulator *rpaA* had a more pronounced effect on metabolism than the deletion of the *kai* genes. The larger impact of the *rpaA* mutation is in agreement with previous transcriptomic analyses and likely relates to a KaiAB1C1-independent function as a transcription factor. Collectively, our data demonstrate an important role of homologs of clock proteins in *Synechocystis* for balanced carbon and nitrogen metabolism during light-to-dark transitions.

## 1 Introduction

Most organisms on Earth experience daily environmental changes. Similar to eukaryotic organisms, photosynthetic cyanobacteria evolved a circadian clock system to respond to predictable fluctuations of light (L) and darkness (D) in a day-night cycle. The freshwater obligate photoautotrophic cyanobacterium *Synechococcus elongatus* PCC 7942 is a model strain used in cyanobacterial circadian research (Cohen and Golden, 2015). Research based on this cyanobacterium established the proteins KaiA, KaiB and KaiC as the core oscillators of the clock system (Ishiura, 1998; Iwasaki, 1999). KaiC performs autokinase, autophosphatase and ATPase functions that are modulated by interaction with KaiA and KaiB. The rhythmic phosphorylation pattern of KaiC functions as a marker of the clock phase and controls rhythmic gene expression. To transfer time information to cellular functions, the oscillator is linked to a two-component regulatory system consisting of the histidine kinase SasA and the DNA-binding response regulator RpaA (Gutu and O’Shea, 2013). During the day, KaiC is progressively phosphorylated. In its highly phosphorylated state, KaiC activates SasA autophosphorylation activity, resulting in the subsequent phosphorylation of RpaA. In *Synechococcus elongatus* PCC 7942, phosphorylated RpaA acts as a repressor of dawn-peaking genes and activator of dusk-peaking genes, and the deletion of *rpaA* leads to the arrest of cells in a dawn-like state (Markson et al., 2013). Although mutations in the KaiABC circadian oscillator do not strongly impair the L/D growth of *Synechococcus elongatus* PCC 7942, the Δ*rpaA* strain is unable to grow in a day/night cycle because it loses viability in D. The described phenotypes of the *Synechococcus rpaA* mutant strain include imbalances in redox regulation and the accumulation of carbon storage compounds (Diamond et al., 2017; Puszynska and O’Shea, 2017).

*Synechocystis* sp. PCC 6803 (hereafter *Synechocystis*) is another frequently used model strain that, in contrast to *Synechococcus elongatus* PCC 7942, can use glucose for mixotrophic and heterotrophic growth and can tolerate high salinities (Rippka et al., 1979; Kirsch et al., 2019). It contains the KaiA, KaiB1 and KaiC1 proteins with high similarity to the *Synechococcus* oscillator proteins. Moreover, the *Synechocystis* genome encodes two additional copies of the *kaiC (kaiC2* and *kaiC3)* and *kaiB* (*kaiB2* and *kaiB3*) genes (Wiegard et al., 2013). Köbler et al. (2018) showed that the homologous SasA-RpaA two-component system of *Synechocystis* interacts only with KaiC1 but not KaiC2 or KaiC3. This specificity of the SasA-RpaA system for KaiC1 suggests that the additional Kai components of *Synechocystis* may use different output elements (**Figure 1**).

**Figure 1.**
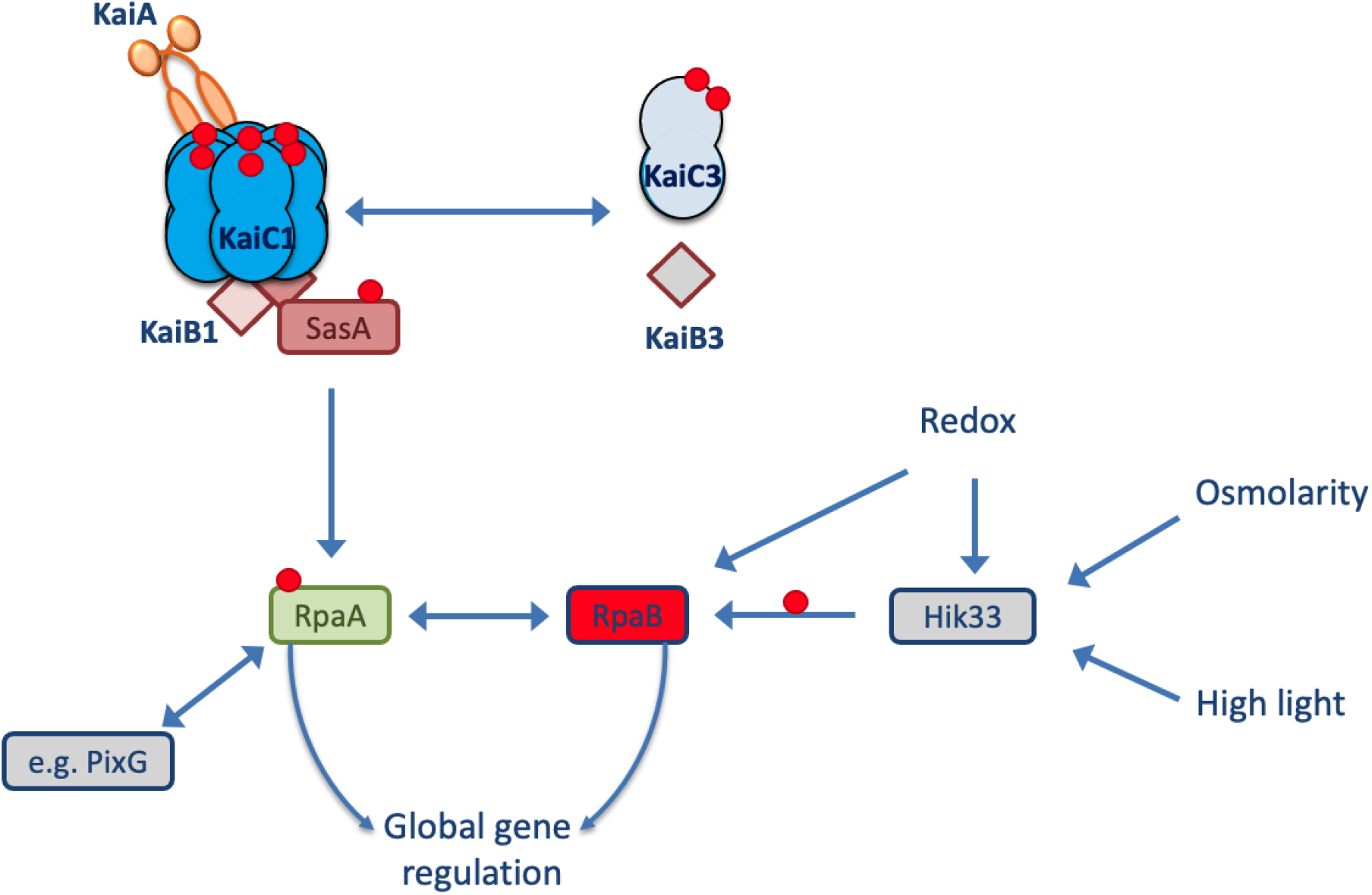
Model of the *Synechocystis* Kai system. In *Synechocystis*, the KaiAB1C1 system is the closest homolog to the well-known circadian clock system from *Synechococcus elongatus* PCC 7942. The output signaling response of KaiC1 relies on the SasA-RpaA two-component system. The transcription factor RpaA controls gene expression in L and D conditions. RpaA can interact with other regulators (e.g Pix and RpaB) and partially targets the same genes as the transcription factor RpaB. The KaiAB1C1 complex is intertwined with the nonstandard KaiB3C3 system in *Synechocystis*, but the direction of signaling between the two systems remains unclear. The red circles represent phosphorylation sites.

True circadian oscillations, which continue after shifting the cells into a constant environment, may not be active in *Synechocystis*, but robust transcriptional rhythmicity in a diurnal light regime has been described (Beck et al., 2014). Diurnal oscillations depend on daily recurring external stimuli sustaining a 24 h period. In *Synechocystis*, the deletion of the *kaiAB1C1* gene cluster and the response regulator gene *rpaA*, caused defects in cell viability in a 12-hour L/12-hour D cycle, especially under mixotrophic conditions. Furthermore, in complete D, neither mutant strain was able to grow heterotrophically, whereas wild-type growth was slow but observable (Dörrich et al., 2014; Köbler et al., 2018). Surprisingly, DNA microarray analyses of both mutant strains demonstrated a considerable difference in their transcriptome responses to a switch from night to day. The overlap in the transcriptional responses between the mutants was only 50% in studies performed under identical conditions with a similar microarray design (Dörrich et al., 2014; Köbler et al., 2018). The overlap in the transcriptional changes contained mRNAs encoding regulatory proteins, such as ribonuclease E (*rne*), phytochrome 1 (*cph1*), and the sigma factor SigE (*sigE*). Furthermore, the Δ*kaiAB1C1* and Δ*rpaA* mutations affected mRNAs of both *gnd* encoding the 6-phosphogluconate dehydrogenase and *talB* encoding the transaldolase. These enzymes are a part of the oxidative pentose phosphate pathway (OPPP), which is activated during the night and converts glucose-6-phosphate (glucose-6P) into pentose-and other sugar phosphates, thereby providing NADPH reduction equivalents (Makowka et al., 2020). In contrast to Gnd, TalB is also active in the Calvin-Benson-Bassham (CBB) cycle during the day. In addition to the common effects of the two mutants, the Δ*rpaA* mutant activates in L the transcription of many genes known to be involved in acclimation to low inorganic carbon (C_i_) concentrations, i.e., bicarbonate and CO_2_. In D, in Δ*rpaA*, but not Δ*kaiAB1C1*, we observed a decreased transcript accumulation of two enzymes (PhaC/PhaE) involved in NADPH-consuming synthesis (via acetoacetyl-CoA reductase PhaB) of the storage compound polyhydroxybutyrate (PHB), which is produced from glycogen under unbalanced nutrient conditions in response to nitrogen starvation (Koch et al., 2019, 2020). Werner et al., (2019) performed very detailed metabolite profiling during a sinusoidal diurnal cycle with a light intensity peak in the middle of the day. These authors found highly concerted oscillations of metabolites across the cycle, verifying that *Synechocystis* performs carbon fixation and protein and nucleotide synthesis during the day. Surprisingly, the authors reported a sharp oscillation in the amount of insoluble carbohydrates at approximately the L-to-D switch. Cyanobacteria store fixed carbon mainly as glycogen, which is then degraded at night using OPPP. Further glycolytic routes, including the Entner-Douderoff (ED) pathway, which branches off the OPPP, and the Embden-Meyerhof-Parnas (EMP) pathway, have been identified (Chen et al., 2016).

The current study aims to investigate how the differential regulation of gene expression in Δ*kaiAB1C1* and Δ*rpaA* mutants impacts *Synechocystis* metabolism. We focused on these two mutants because they showed a growth phenotype in a L-D cycle in contrast to a Δ*kaiC3* strain which had no growth defect under these conditions and a *kaiB2C2* mutation which turned out to be lethal (Dörrich et al., 2014). For this purpose, targeted and untargeted metabolic profiling was applied to reveal the changes in primary metabolites in the middle of the D and L phases. The same time points were investigated in a previous transcriptome analysis of the Δ*kaiAB1C1* and Δ*rpaA* mutants. The identified differences in the metabolic responses clearly indicate that mainly the lack of the RpaA regulator impacts primary metabolism and the metabolic switch between L and D growth.

## 2 Material and Methods

### 2.1 Cyanobacterial strains, culture and sampling conditions

For standard cultivation, the *Synechocystis* sp. PCC 6803 wild-type (PCC-M, resequenced, Trautmann et al., 2012) and mutant strains were grown photoautotrophically under constant illumination with 75 μmol photons m^−2^ s^−1^ white light at 30°C in BG11 medium (Rippka et al., 1979) supplemented with 10 mM TES buffer (pH 8) in flasks. For the L/D experiments, the precultures were grown in constant light and diluted to OD_750 nm_ 0.4-0.6 before transfer to alternating 12-hour L/12-hour D cycles with 75 μmol photons m^−2^ s^−1^ during the L phase. Depending on the experimental setup, air was supplemented with CO_2_ (1%, v/v), and illumination was increased to 150 μmol photons m^−2^ s^−1^ after the liquid cultures reached an OD_750 nm_ of ~1. The wild-type and mutant strains were cultured in triplicate. The samples were collected in the middle of the first D phase and the following L phase 5.5 h after each transition in the exponential growth phase if not described otherwise.

### 2.2 RNA isolation and Northern blot hybridization

The RNA isolation was performed following Pinto et al., (2009). For sampling, 20 ml of culture were filtered through a Supor-800 membrane filter (0.8 mm, Pall), vortexed in 1.6 ml PGTX for 30 s and immediately frozen in liquid nitrogen. The samples were stored at −80°C. After thawing the samples on ice, the samples were heated at 95°C for 5 min. The lysis was supported by vortexing during the incubation. After cooling the samples on ice for 5 min, the samples were incubated at room temperature (RT) for 10 min. Then, 200 μl of cold BCP were added, vortexed for 30 s and stored at RT for 15 min. Next, the phases were separated by centrifugation at 3100 ×g for 15 min at 4°C. The aqueous phase was transferred into a new reaction tube, and the reaction was repeated once more with 1 volume of cold BCP. The samples were centrifuged at 21130 ×g for 15 min at 4°C, followed by precipitation with 1 volume of isopropanol and storage of the sample overnight at −20°C. The samples were centrifuged at 21130 ×g for 1 h at 4°C, the resulting pellet was washed with 75% EtOH, and centrifugation was repeated for 10 min. The pellet was air-dried for 15 min and resuspended in 40 μl ddH_2_O.

To separate the RNA, 10 μg were used for electrophoresis on a 1.5% denaturing agarose-formaldehyde gel. The gel was blotted onto a Roti-Nylon plus membrane (Roth). The DNA fragment used for probe generation was amplified from wild-type DNA using the oligonucleotides sbtA-fw (5’ CAATCCCCATAAATTTCAACCAAGGAGAC 3’) and sbtA-rev (5’ TAATACGACTCACTATAGGGAGAGCCAAGGGCGGCAATAACCATC 3’). The probe was produced with a Rediprime II labeling kit (GE Healthcare). Hybridization was performed as described by Dienst et al. (2008), and the signals were detected using a Typhoon FLA 9500 (GE Healthcare).

### 2.3 Immunoblot analysis of RubisCO

The wild-type and mutant cultures of *Synechocystis* were diluted to OD_750 nm_ 0.6 and cultivated in an L/D cycle. 5 ml of WT and mutant cultures of *Synechocystis* were harvested by centrifugation at 4000 ×g for 2 min at 4°C. After the pellet was resuspended in PBS buffer (8.6 mM Na_2_PO_4_, 1.8 mM KH_2_PO_4_, 137 mM NaCl, 2.7 mM KCl, pH 7.5), the cells were disrupted in a cell mill using glass beads. The crude extract was obtained by centrifugation at 500 ×g for 5 min at 4°C, and different protein amounts of the crude extract were subjected to SDS-PAGE. The immunodetection was performed using anti-RbcL (Agrisera, Sweden) and secondary anti-rabbit (Thermo Fisher Scientific Inc., USA) antibodies.

### 2.4 Quantification of the PHB content

The intracellular PHB content was measured as described by Taroncher-Oldenburg et al. (2000). The wild-type and mutant cultures were adjusted to OD_750 nm_ 0.2 and then grown in L/D cycles supplemented with CO_2_ as described above. The samples were collected in the 7^th^ L/D cycle 1 h before the L or D transitions. Approximately 15-35 ml of cell culture were harvested by centrifugation (4000 ×g, 10 min, 4°C), washed once with H_2_O and dried overnight at 85°C. To break the cells and convert PHB, the pellet was boiled in 1 ml concentrated H_2_SO_4_ for 1 h. Then, 100 μl were transferred to 900 μl of 0.014 M H_2_SO_4_. To pellet the cell debris, the sample was centrifuged at 17000 ×g for 5 min before 500 μl of the supernatant were diluted in 500 μl of 0.014 M H_2_SO_4_. The centrifugation was repeated, and the supernatant was analyzed via Acquity UPLC BEH (WATERS) with a reversed-phase C18 column (Eurosphere II, 100 by 2.1 mm, Knauer). The separation was performed using increasing concentrations of acetonitrile in 0.1% formic acid. Commercially available crotonic acid was used as a standard in parallel with a conversion ratio of 0.893 (Koch et al., 2019).

### 2.5 Nontargeted metabolome profiling analysis by gas chromatography-mass spectrometry (GC-MS)

The metabolome profiling of the metabolite fraction enriched with primary metabolites was performed as previously described (Kopka et al., 2017). 20 ml of the cultures were sampled by fast filtration onto 47 mm diameter glass microfiber filters of a 1.2 μm pore size (GE Healthcare, Little Chalfont, England), followed by immediate shock freezing. The complete process was less than 30 s. Samples equivalent to approximately 20 ml of OD_750 nm_ = 0.7-1.0 cultures were extracted for 15 min at 37°C by 1 ml methanol:chloroform:diethylamine 2.5:1.0:1.0 (v/v/v). The extraction mixture contained 0.02 mgml^−1 13^C_6_-sorbitol, which was used as an internal standard (Erban et al., 2020). A polar liquid phase was obtained by adding 400 μl distilled water and centrifuging for 5 min at 14,000 rpm in an Eppendorf 5417 microcentrifuge. The top liquid phase, ~700 μl, was dried by vacuum centrifugation overnight in 1.5-mL microcentrifugation tubes and stored at −20°C.

The chemical derivatization for the GC analysis, GC–MS-based metabolite profiling and retention index standardization were performed as described by Erban and coauthors (2020) using electron impact ionization time-of-flight mass spectrometry (GC/EI-TOF–MS). The GC/EI-TOF–MS system was an Agilent 6890N24 (Agilent Technologies, Waldbronn, Germany) gas chromatograph hyphenated to a LECO Pegasus III time-of-flight mass spectrometer (LECO Instrumente GmbH, Mönchengladbach, Germany). The chromatograms were acquired, and the baseline was corrected by ChromaTOF software (LECO Instrumente GmbH, Mönchengladbach, Germany). The metabolites were annotated by manual supervision using TagFinder software (Luedemann et al., 2008), NIST2017 version 2.3^1^ and the mass spectral and retention time index (RI) reference data of the Golm Metabolome Database (Kopka et al., 2005). GC/EI-TOF–MS was normalized to the internal standard ^13^C_6_-sorbitol, OD_750 nm_ of the culture and volume of the sample.

### 2.6 Targeted metabolome analysis by liquid chromatography-mass spectrometry (LC-MS)

The sampling of the cells was performed precisely as described in the previous paragraph. Low molecular mass compounds were extracted from the cells with 2 ml of ethanol (80%, HPLC grade, Roth, Germany) at 65°C for 2 h. One microgram of carnitine was added to each sample as an internal standard to correct for losses during the extraction and sample preparation. After centrifugation, the supernatants were collected and freeze-dried. The dry extracts were dissolved in 1 ml MS-grade water and filtered through 0.2 μm filters (Omnifix®-F, Braun, Germany). The cleared supernatants were analyzed using a high-performance liquid chromatograph mass spectrometer system (LCMS-8050, Shimadzu, Japan). In brief, 1 μl of each extract was separated on a pentafluorophenylpropyl (PFPP) column (Supelco Discovery HS FS, 3 μm, 150 × 2.1 mm) with a mobile phase containing 0.1% formic acid. The compounds were eluted at a rate of 0.25 ml min-1 using the following gradient: 1 min 0.1% formic acid, 95% distilled water, 5% acetonitrile, within 15 min linear gradient to 0.1% formic acid, 5% distilled water, 95% acetonitrile, 10 min 0.1% formic acid, 5% distilled water, and 95% acetonitrile. Aliquots were continuously injected into the MS/MS part and ionized via electrospray ionization (ESI). The compounds were identified and quantified using the multiple reaction monitoring (MRM) values given in the LC-MS/MS method package and the LabSolutions software package (Shimadzu, Japan). Defined standard mixes of amino acids and organic acids (usually 1 ng of the specific metabolite was injected) were analyzed in the same run to calculate the absolute contents per sample. Then, the metabolite levels were normalized to the detected amount of the internal standard carnitine, sample volume and OD_750 nm_.

### 2.7 NAD(P) and NAD(P)H analyses

NAD(P) and NAD(P)H were extracted from approximately 8 ml of OD_750 nm_ = 0.7-1.0 cultures sampled onto 47 mm diameter glass microfiber filters of a 1.2 μm pore size (GE Healthcare, Little Chalfont, England), followed by immediate shock freezing. The quantification of NAD, NADP and NADPH was performed as previously described using enzymatic cycling assays (Zhang et al., 2020). The samples used for the NAD and NADP analyses were extracted by 500 μL extraction solution containing 0.1 M HClO_4_ in 50% ethanol; the NADPH extraction was performed by 0.1 M KOH in 50% ethanol (Tamoi et al., 2005). The samples were thoroughly vortexed with intermittent sonication in an ice bath 3 times for 45 sec each, followed by a final incubation on ice for 10 min. After centrifugation for 10 min at 14,000 rpm at 4°C, 100 μL of extract were transferred to a microcentrifuge vial and heat inactivated at 95°C for 2 min. Then, the samples were cooled on ice and neutralized either by 100 μL of 0.1 M KOH in 0.2 M Tris HCl pH 8.4 for the NAD and NADP quantitation or 100 μL of 0.1 HClO_4_ in 0.2 M Tris HCl pH 8.4 for the NADPH analyses. The quantitation was performed using 25 or 40 μL volumes of the neutralized extracts used for the previously described cycling assays (Zhang et al., 2020). Care was taken to adjust the samples and quantitative calibration standards to identical final ethanol concentrations (Supplemental Table S3). In this study, the quantities of NAD, NADP and NADPH in the biological samples were sufficient for the quantitation. NADH quantitation was attempted but failed due to low concentrations in the biological samples. The analyses were biologically replicated using 3 independent cultures. The cultures were probed by 1 (NADPH) or 2 (NAD and NADP) analytical repeats, i.e., parallel samplings from the same culture. The quantitative data of the analytical repeats were averaged prior to the statistical analysis of the 3 independent biological replications.

### 2.8 Metabolomic data analysis

All metabolomics data analyses, the hierarchical cluster analysis (HCA) by Pearson’s correlation and average linkage, the principal component analysis (PCA), the two factorial analyses of variance (ANOVA), the statistical testing, e.g., Tukey tests, and the data visualization, were performed using RStudio statistical programming^2^ and the program packages “impute”, “multcompView”, “ggplot2”, “eulerr”, and “https://github.com/MSeidelFed/RandodiStats_package.git”. The bootstrapping analysis of the HCA tree was performed using the “pvclust” R package, and the node confidence was evaluated by “approximately unbiased” and “ordinary” bootstrap probabilities.

The relative quantification data from the GC-MS profiling analyses and quantitative data from the LC-MS analyses were maximum scaled per metabolite prior to further analyses. For the global data analyses, the missing values in the data set were replaced by the k-nearest neighbors algorithm (k-NN) and autoscaled. For the statistical data analyses, the data matrices were log_10_-transformed after the missing data replacement by a small value, i.e., 10^−9^. The statistical analyses were performed using the log-transformed data. The relative concentration changes are reported as fold changes (FCs). The FCs of the metabolites that were detectable under one condition and undetectable under the compared condition, e.g., FCs comparing L to D responses or FCs comparing mutant to wild type (**Supplemental Table S1-S2**), are reported. The significance analyses were replaced by presence and absence calls, and the FCs were scored as either significant increases or decreases.

For the analyses comparing the data to previous studies, the data were merged according to the technology platform, namely, GC-MS profiling and LC-MS quantification. Merging across technology platforms was only performed if data from the same technology platform were not available. The replicate data were averaged per study maximum-scaled, and the missing values were replaced by the k-NN method. The final data matrices were log transformed and autoscaled prior to the subsequent PCA.

## 3 Results

### 3.1 CO_2_ availability can compensate for the growth defects of Δ*kaiAB1C1* and Δ*rpaA* mutants in L/D cycles

Previous analyses revealed the essential role of the clock homologs KaiAB1C1 and RpaA in the viability of *Synechocystis* in L/D cycles and in the D under heterotrophic conditions (Dörrich et al., 2014; Köbler et al., 2018). Although the phenotypes of the corresponding mutants were very similar, microarray analyses revealed substantial differences in gene expression between these strains, such as an accelerated C_i_ response that was detected only in the Δ*rpaA* mutant (Köbler et al., 2018). To determine whether C_i_-responsive genes are still upregulated in Δ*rpaA* even under high CO_2_ conditions, we analyzed the expression of *sbtA*, which encodes a sodium-dependent bicarbonate transporter. Using Northern blot analysis, we did not detect a difference in the *sbtA* mRNA levels between the wild-type and mutant strains under elevated CO_2_ concentrations, suggesting that C_i_-responsive genes are not a direct target of the RpaA transcription factor (**Supplementary Figure S1**). Furthermore, we examined the response of the mutant strains to CO_2_ availability by growing cells under low (ambient) and high CO_2_ (1%) conditions in liquid cultures (**Figure 2**). Similar to previous studies in which viability assays were performed using agar plates, the Δ*kaiAB1C1* and Δ*rpaA* strains showed a pronounced growth defect in L/D cycles (**Figure 2A**). The curves also revealed that growth arrest mainly appeared when the mutant cultures left the exponential growth phase (**Figure 2A**). As an organic carbon source, glucose had an additional inhibitory effect on the viability of the mutant cells (Dörrich et al., 2014; Köbler et al., 2018), whereas an increase in CO_2_ availability could largely align the growth of the mutant cultures with the wild-type cultures in an L/D cycle (**Figure 2B**). Our previous viability analyses under mixotrophic/heterotrophic conditions and the growth assays under elevated CO_2_ availability in this study indicate that defects in the *Synechocystis* Kai system led to an imbalance in carbon acquisition and changes in central carbon metabolism.

**Figure 2.**
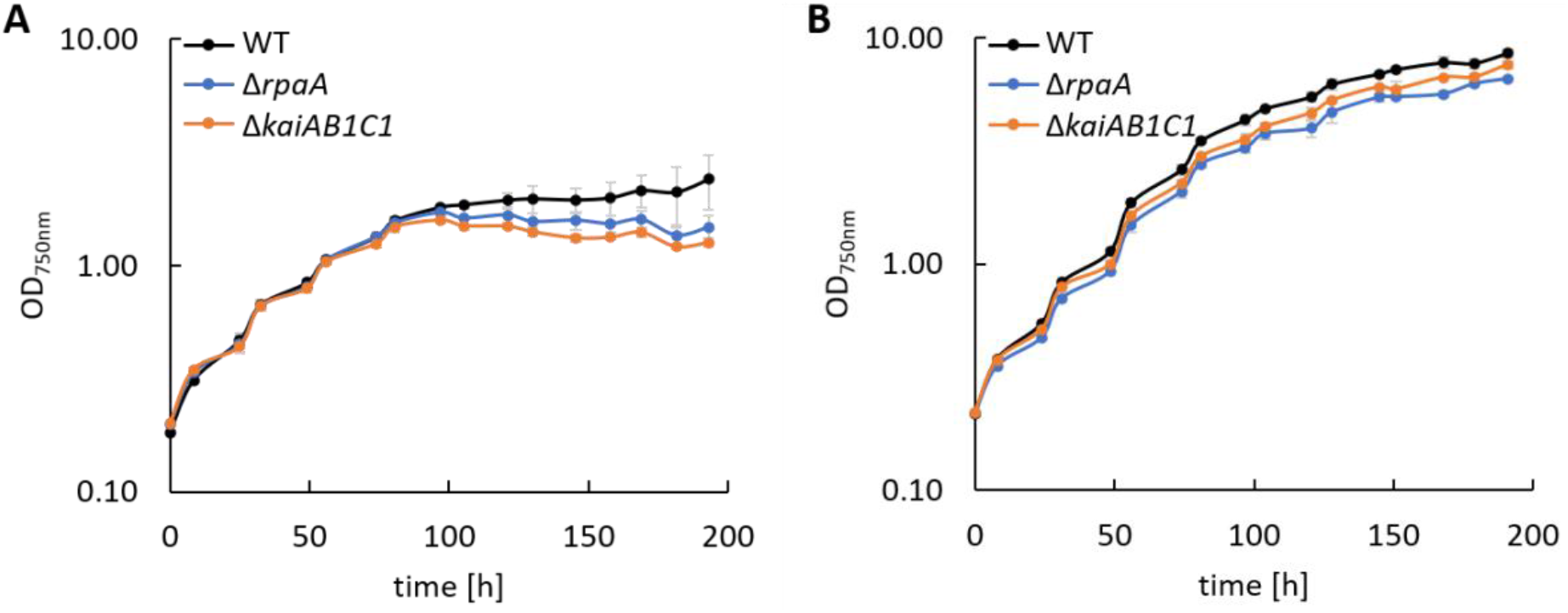
Growth of *Synechocystis* mutants. Wild type (WT), Δ*rpaA* and Δ*kaiAB1C1* deletion mutants were grown in a 12-hour L/12-hour D cycle with ambient air **(A)** and 1% CO_2_ **(B)**. The light intensity during cultivation was 75 μmol photons m^−2^ s^-1,^ which was increased to 150 μmol photons m^−2^ s^−1^ in **B** after the cultures reached an OD_750 nm_ of ~1. Each point represents the mean of three biological replicates (± standard deviation).

### 3.2 PHB accumulation is reduced in *Synechocystis* 6803 Δ*kaiAB1C1* and Δ*rpaA* mutants

In addition to the altered C_i_ acclimation response, the expression of two genes encoding polyhydroxybutyrate (PHB) synthase subunits (*phaC* and *phaE*) was highly decreased in the Δ*rpaA* strain in D (Köbler et al., 2018). A similar tendency in *phaCE* transcript accumulation was also detected in a microarray analysis of the Δ*kaiAB1C1* strain; this tendency was less pronounced and did not match our significance criteria (Dörrich et al., 2014). In the previous microarray analyses, the cells were collected during the exponential growth phase and were grown under low CO_2_ conditions, where PHB synthesis is very low, and the amount of this storage compound was not measurable. Therefore, we evaluated the impact of KaiAB1C1 and RpaA on PHB production under high CO_2_ conditions during the stationary growth phase. Under these conditions, the Δ*kaiAB1C1* and Δ*rpaA* strains did not show growth arrest in an L/D cycle and exhibited only a slightly lower growth rate than the wild type (**Figure 2B**). Consistent with the lowered *pha* gene expression, both strains showed a strong reduction in PHB accumulation in the D and L phases, suggesting that the high CO_2_ availability was unable to compensate for the defect in PHB production (**Figure 3**). PHB synthesis depends on many factors, such as the C:N ratio and nutrient starvation. A complex relationship links PHB synthesis and utilization to central metabolic pathways (Ciebiada et al., 2020; Koch et al., 2020). Therefore, the transcriptomic changes might not fully represent the role of KaiAB1C1 and RpaA in cellular night- and daytime functions, which led us to analyze the metabolic changes in the two corresponding mutants.

**Figure 3.**
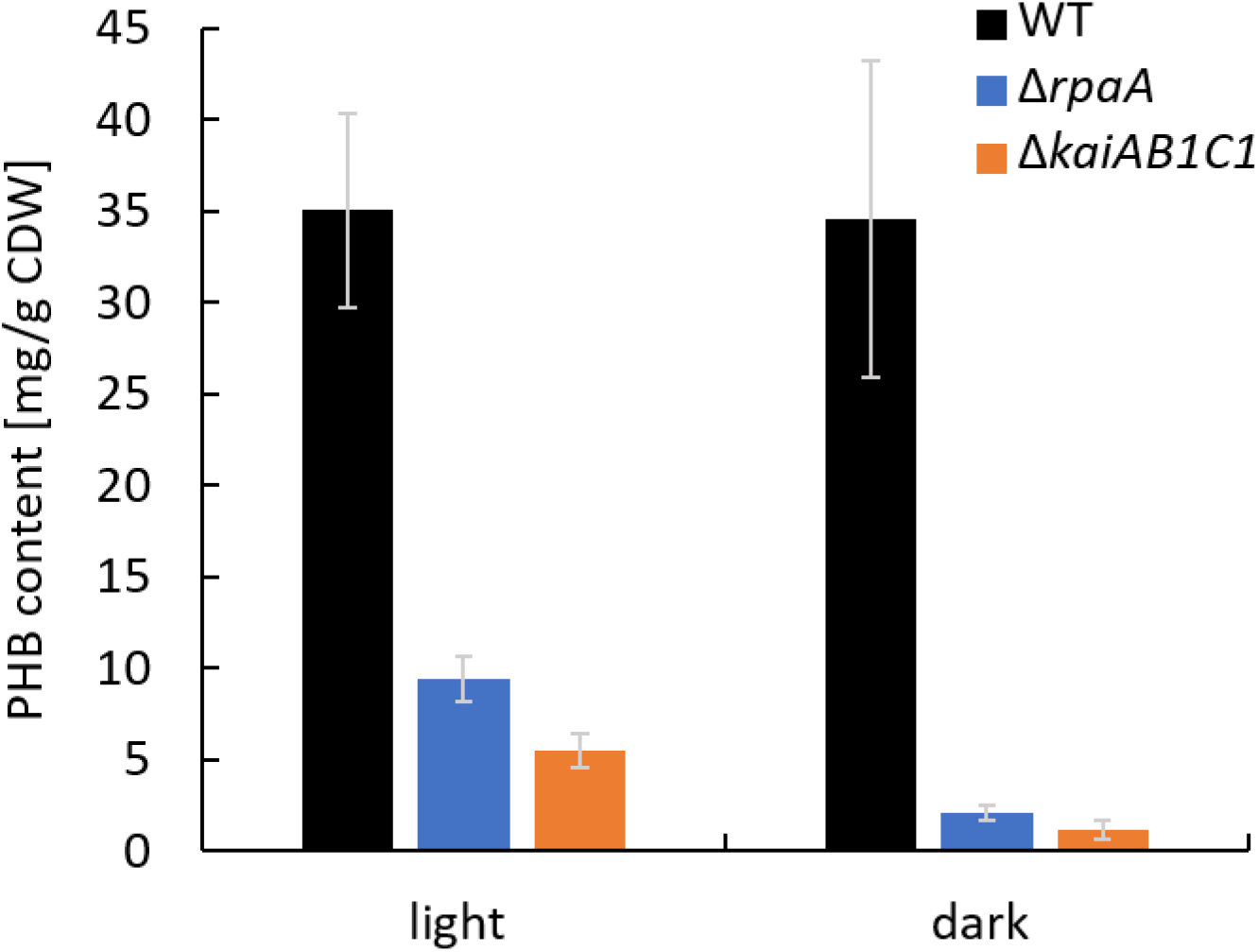
The Δ*rpaA* and Δ*kaiAB1C1* strains show lower PHB accumulation at high CO_2_ concentrations. PHB was quantified in cells cultivated in a 12-hour L/12-hour D cycle and 1% CO_2_. The samples used for the PHB measurements were collected in the stationary phase at an OD_750 nm_ of ~7 in the L and D phases of the 7^th^ L/D cycle one hour before switching. Each bar displays the mean of three technical replicates (± standard deviation).

### 3.3 Global metabolic analysis of the Δ*rpaA* and Δ*kaiAB1C1* mutants after L/D entrainment

The previous investigations and data presented above suggest that the KaiAB1C1 proteins along with the RpaA regulator impact primary metabolism in *Synechocystis* and growth specifically under low C_i_ conditions in an L/D cycle (**Figure 2A**). Therefore, the metabolome of the two mutants was analyzed after an L-to-D transition under ambient CO_2_ supply. Independent, exponentially grown precultures in ambient air were adjusted to similar optical densities of approximately 0.6 and sampled either in the middle of the first D phase at 5.5 h of the diurnal cycle or in the middle of the following L phase at 17.5 h (**Figure 4A**). The mutants and wild type exhibited similar growth rates during this precultivation under continuous L (**Supplementary Figure S2**). The time points were selected to avoid the expected rapid changes in metabolism directly after illumination shifts (Werner et al., 2019), to minimize the secondary effects of the growth arrest of the mutants in L/D cycles and to correlate the metabolic data of this study to previously reported transcriptomic data (Dörrich et al., 2014; Köbler et al., 2018) and to metabolic data published for *Synechococcus elongatus* PCC 7942 (Diamond et al., 2017).

**Figure 4.**
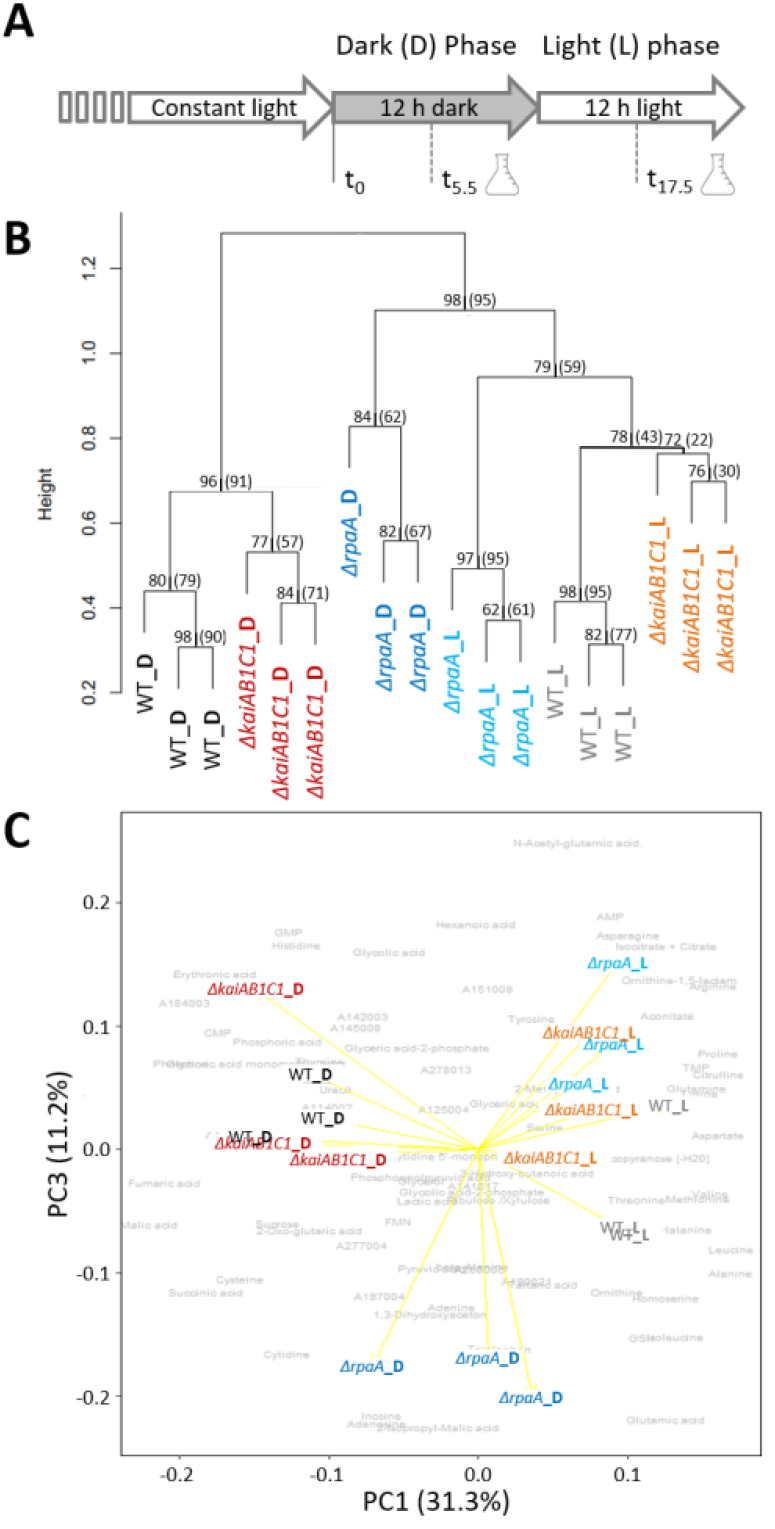
Global analysis of the metabolic L/D switch in the Δ*kaiAB1C1* and Δ*rpaA* mutants. **(A)** Design of the two-factorial entrainment experiment. Wild-type and mutant cells, i.e., the first experimental factor (genotype), were precultivated under continuous light, divided into independent cultures of OD_750 nm_ = 0.6 in ambient air, and sampled in the middle of the first night (D phase) and first day (L phase) after entrainment, i.e., the second experimental factor (illumination). Three independent replicate cultures in each phase were analyzed by a quantitative targeted LC-MS analysis and nontargeted GC-MS metabolome profiling of the primary metabolome. **(B)** Hierarchical cluster analysis (HCA) of mutant and wild-type metabolomes. Annotated LC- and GC-MS metabolite data separately maximum scaled and then combined, log-transformed, autoscaled, and clustered using Pearson’s correlation distance metric and average linkage. HCA includes a scale of the node height and bootstrap-support analysis by 1000 iterations. The values represent the ‘approximately unbiased’ bootstrap probability, and in brackets, the conventional bootstrap probability obtained using the ‘pvclust’ R package is shown. Values of 100 represent the highest node probability. **(C)** Principal component analysis (PCA) of the same data. PC1 and PC3 demonstrate the variance contribution of illumination (PC1) and the Δ*rpaA* samples collected in the D phase (PC3) to the metabolomics data set. Yellow arrows represent the score biplot of the samples. The gray underlay indicates the loading contributions of the metabolites. A corresponding supplement contains a biplot of PC1 and PC2 and a scree plot of the variance represented by the top 10 PCs (**Supplemental Figure S3**).

We analyzed 80 metabolites from central metabolism by relative quantification of a metabolite fraction enriched for primary metabolites via gas chromatography-mass spectrometry (GC-MS)-based profiling or absolute quantification via liquid chromatography-mass spectrometry (LC-MS) (**Supplemental Table S1**). In addition to the identified metabolite information, the nontargeted metabolite profiling yielded a set of yet unidentified metabolites and mass features that contributed to the global analyses of the metabolic phenotypes in this study. The hierarchical cluster and principal component analyses (PCA) indicated that the switch between D and L was the major experimental factor affecting the metabolite phenotypes (**Figure 4B**) with a contribution of 31.3% to the total variance according to PC1 (**Figure 4C**, **Supplemental Figure S3**). The wild type and two mutants clustered with high bootstrapping confidence of their independent replicate profiles (**Figure 4B**). The metabolic profiles of the Δ*kaiAB1C1* and Δ*rpaA* mutants differed from those of the wild type in the D and L phases (**Figure 4B-C**). However, the metabolic phenotype of the Δ*rpaA* mutant differed more, especially in D, when the deviations were stronger and amounted to a distinct PC3. PC3 differentiated Δ*rpaA* D metabolism from all other sample profiles by 11.2% of the total variance (**Figure 4C**). The D metabolism of Δ*rpaA* appeared to be between that of the wild type D and L states (**Figure 4C**) with a tendency towards wild type in L (**Figure 4B**).

Molecular studies have suggested that the Δ*rpaA* mutant of *Synechococcus elongatus* PCC 7942 may be arrested in a dawn-like expression state (Markson et al., 2013). In wild-type *Synechocystis* cells, a set of metabolites accumulates rapidly following the onset of illumination after dawn and declines in the course of the L phase, which may indicate a morning state of metabolism (Werner et al., 2019). These “morning metabolites” include malate, fumarate, amino acids, serine, valine, isoleucine, glutamine, aspartate, arginine, and nucleotide-related metabolites, such as cytidine, cytidine monophosphate (CMP), cytidine triphosphate (CTP), uridine monophosphate (UMP), guanosine monophosphate (GMP), guanosine triphosphate (GTP), hypoxanthine, inosine, and inosine monophosphate (IMP) (Werner et al., 2019). Although our study investigated the metabolome in the middle of the day, two “morning metabolites”, namely, aspartate and serine, significantly accumulated (Tukey test, *P* < 0.05) in the illuminated Δ*rpaA* mutant of *Synechocystis*, whereas many more metabolites shifted to higher values in the wild-type cells. In contrast, isoleucine and valine significantly accumulated during D in Δ*rpaA* (**Figure 5**). For a global comparison of our metabolite profiles and a previous diurnal characterization of *Synechocystis* central metabolism (Werner et al., 2019), we aligned the 47 commonly monitored metabolites. The PCA of the combined and separately scaled data sets revealed PC1, which had a high percentage of shared total variance, i.e., 45.8%. PC1 separated the wild-type L metabolomes from the D metabolomes of the two studies in a nonlinear manner (**Supplemental Figure S4**). Colinearity indicated high agreement in the L/D-induced metabolome changes between the two studies. However, PC2, which accounted for 16.8% of the total variance, rendered a more precise alignment, and a comparison of the sampled time points between the two studies is impossible. Further comparative meta-analysis was abandoned because the studies clearly used different growth conditions. For example, high peaking light intensity at 1600 μmol photons m^−2^ s^−1^ and high 5% CO_2_ were applied in a previous study (Werner et al., 2019), whereas low light of 50 μmol photons m^−2^ s^−1^ and ambient CO_2_ were applied in this study.

**Figure 5.**
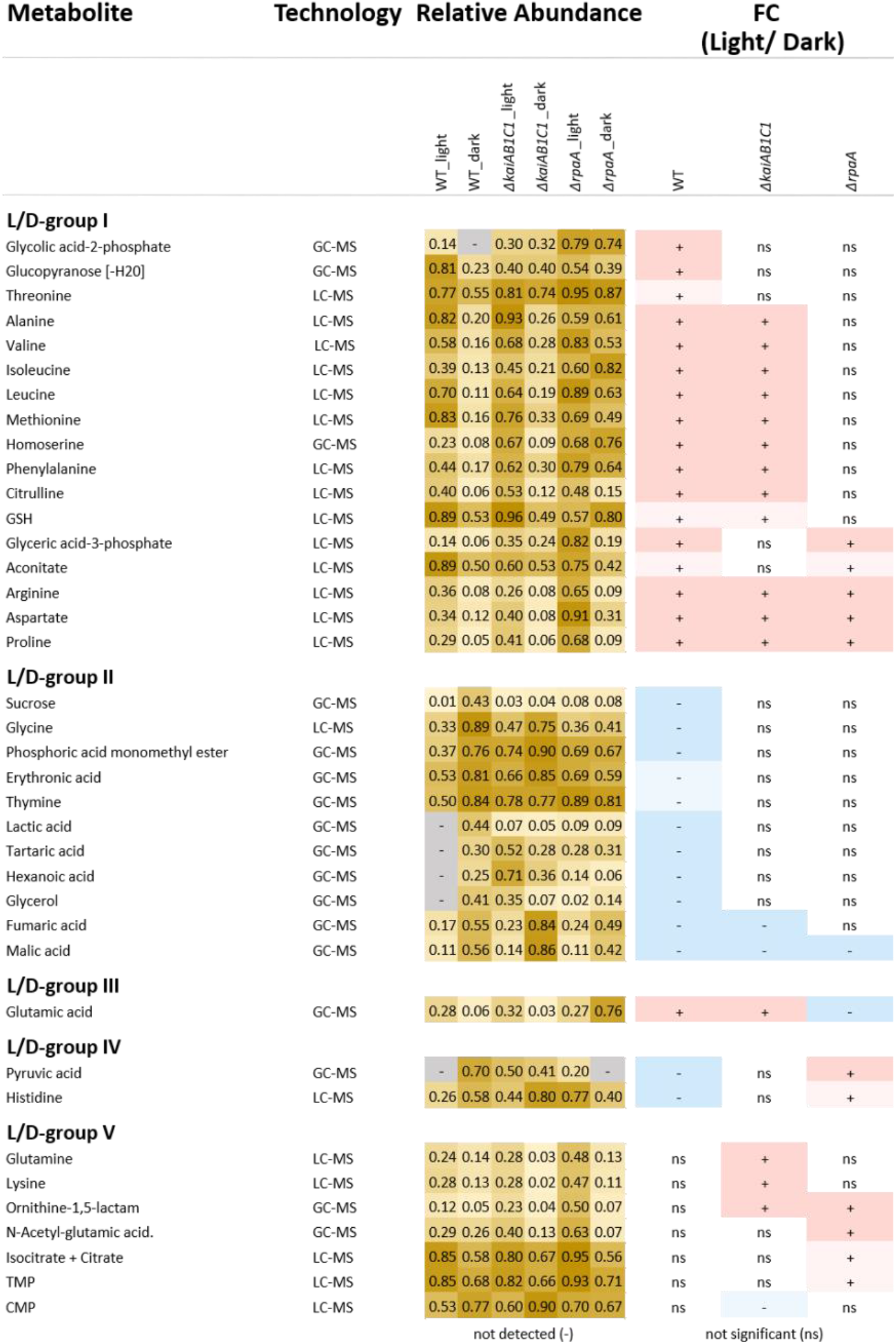
Relative abundances and fold changes (FC) of metabolite concentrations in the L phase compared to the D phase. The figure includes identified metabolites that exhibited significant changes in L metabolism compared to D metabolism (Tukey test P < 0.05). Fold changes are color-coded as follows: significant increases, red (=2-FC), light red (<2-FC), significant decreases, blue (<0.5-FC), and light blue (=0.5-FC); the complete numerical data are provided in **Supplemental Table S1.** The grouping of the metabolites into the L/D response groups (L/D groups) was performed according to the changes in the wild type (L/D group I-II) and considered inverted (L/D group III-IV) changes in the mutants or emergent changes (L/D group V). Heat map: light yellow, low abundance; dark yellow, high abundance; gray, undetectable.

A global correlation analysis of the metabolites was performed using all mutant and wild-type data in the current study. We present this analysis as a Pearson’s correlation matrix that analyzes the relative concentration changes in all monitored metabolites (**Supplemental Figure S5**). This analysis indicates a central group of highly correlated metabolites that contained metabolites of lower glycolysis, 3-phosphoglycerate (3PGA), 2-phosphoglycerate (2PGA), phosphoenolpyruvate (PEP), and of photorespiration, 2-phosphoglycolate (2PG), serine and glycerate. Two additional large groups contained metabolites that were inversely correlated between the groups but partially formed highly correlated subsets within each group. The first group comprised most amino acids and the oxidative (C6-C5) branch of tricarboxylic acid (TCA) metabolism. The second group contained predominantly metabolites of nucleic acid metabolism and the reductive (C4) branch of TCA metabolism (**Supplemental Figure S5**).

### 3.4 Δ*kaiAB1C1* and Δ*rpaA* mutations attenuate metabolic L/D responses

Our study revealed that 46 metabolites, i.e., more than half of all monitored compounds, significantly (Tukey test, *P* < 0.05) differed between the mutants and wild type. A large overlap of 21 metabolic changes during the L/D cycle was observed between the two mutants (**Figure 6**). This finding supported the hypothesis that the KaiAB1C1 system contributes to the diurnal control of central metabolism mostly via RpaA-mediated output signaling (**Figure 1**). Most of the common metabolic defects were apparent in both phases (12), except for one defect, which was limited to the L phase (9). Deviations specific to only one mutant suggested a small RpaA-independent metabolic signaling component in the KaiAB1C1 system, with three metabolic changes in D, one metabolic change in L and one metabolic change in both. In contrast, the KaiAB1C1-independent changes in the Δ*rpaA* mutant (20) exhibited a preference for RpaA-specific metabolic defects in D (10) (**Figure 6**), which is consistent with the closer clustering of Δ*rpaA* D with the L samples of all strains (see **Figure 4**). The larger impact of the RpaA deficiency may be explained by its functionality as a transcription factor with partly retained functions in the absence of Kai-based control and the KaiAB1C1-independent additional functions of RpaA that are established via protein-protein interactions (Köbler et al., 2018). For example, RpaA interacts with RpaB, which is the transcriptional regulator responding to environmental changes, including L/D transitions, via concomitant redox regulation (Riediger et al., 2019) (**Figure 1**).

**Figure 6.**
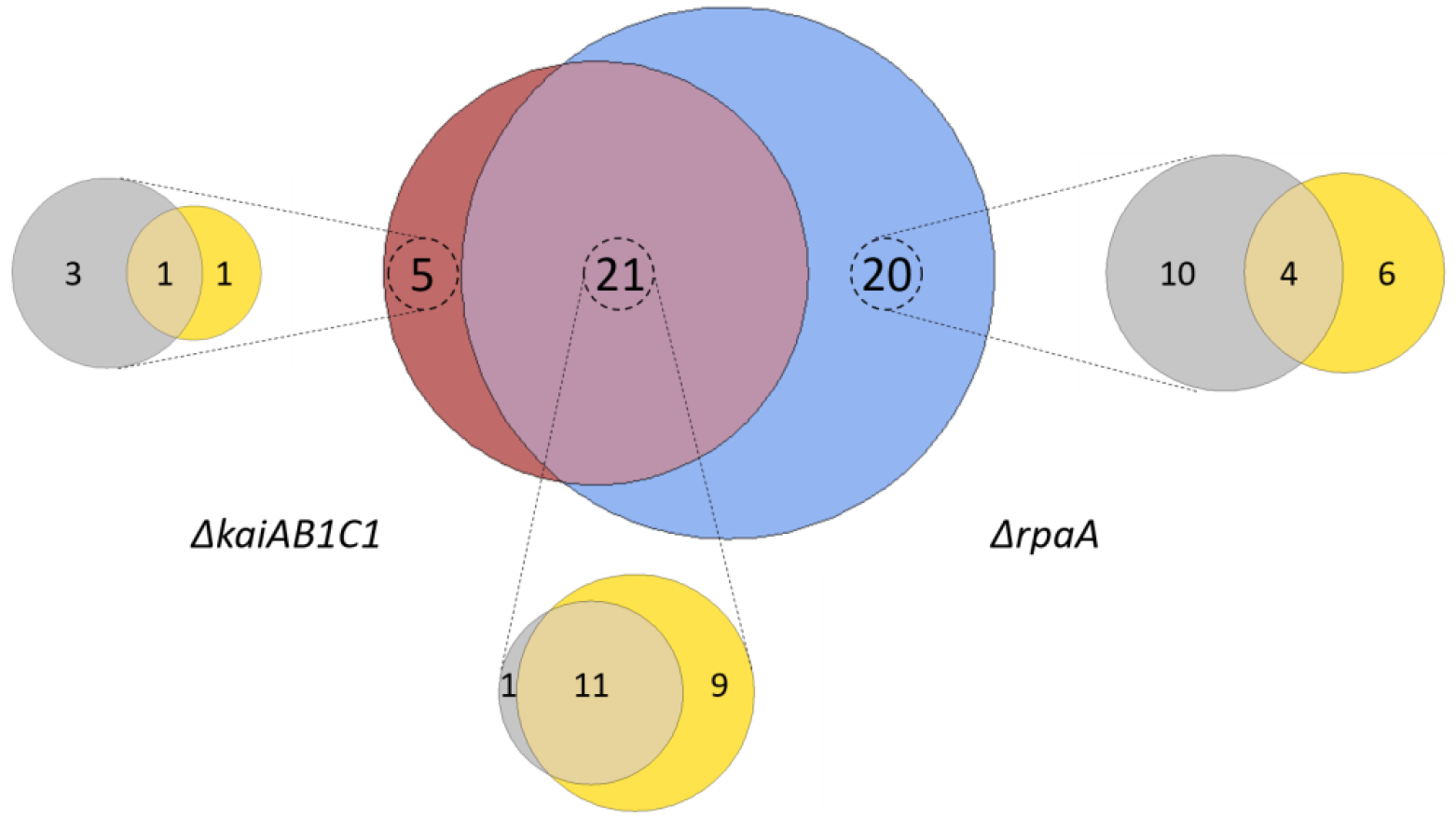
Summary of the metabolic changes in the mutants of *Synechocystis*. Venn diagram analysis of all significant (Tukey test, *P* < 0.05) metabolic changes among the set of identified and yet unidentified metabolites. Intersections show significant changes in the metabolite pools shared between the Δ*kaiAB1C1* (brown) and Δ*rpaA* (blue) mutants. Zoom-ins show the contribution of the two diurnal phases to the respective sets, D phase (gray) and L phase (yellow). Note the large overlap between the mutants and the more frequent specific responses of the Δ*rpaA* cells.

The set of 46 mutant-dependent metabolites included unidentified compounds and metabolites with small but significant pool size changes (**Supplemental Table S1**). To focus our data analysis, for the following analysis, we selected only identified metabolites that were significantly (Tukey test, *P* ≤ 0.05) responsive to the L/D change (**Figure 5**). From this subset, we distinguished five groups of L/D-responsive metabolites (L/D groups) according to their relative changes in the wild type compared to the mutant defects. L/D group I contained 17 metabolites (mostly amino acids) that accumulated in the wild type in L compared to D (**Figure 5**). A second set of 11 metabolites, i.e., L/D group II, exhibited inverse changes in the wild type compared to group I and contained metabolites, e.g., organic acids, that decreased in L compared to D. Only a small set of metabolites from these two groups, namely, arginine, aspartate, proline (L/D group I) and malate (L/D group II), maintained L/D responsiveness in the mutants. KaiAB1C1 and especially the RpaA deficiency attenuated the L/D responsiveness of a large fraction of L/D-responsive metabolites in both groups. Many attenuated changes were shared between the Δ*rpaA* and Δ*kaiAB1C1* mutants. However, the Δ*rpaA* mutant clearly had more metabolites with attenuated L/D responsiveness (**Figure 5**).

Furthermore, we discovered additional but rarer metabolite response patterns supporting the hypothesis that central metabolic responses to a switch between photoautotrophy and dark-induced heterotrophy are all but passive. These metabolic responses are enhanced and modulated by KaiAB1C1 and/or RpaA. Glutamate (L/D group III) represented one of these rarer patterns as it accumulated in the wild type in the L phase and had an inverted L/D response only in the Δ*rpaA* mutant. Similarly, pyruvate and histidine (L/D group IV) were depleted in the wild type in the L and accumulated in the Δ*rpaA* mutant (**Figure 5**). Some metabolites (L/D-group V), e.g., N-acetyl-glutamate, glutamine, ornithine-1,5-lactam, lysine, citrate/isocitrate, TMP and CMP, even gained significant L/D responsiveness in one or both mutants (**Figure 5**). Further central metabolites, e.g., 2PGA and PEP, exhibited response patterns similar to those of L/D group IV but had only small effect sizes and failed to pass the significance threshold (**Supplemental Table S1**).

### 3.5 Δ*kaiAB1C1* and Δ*rpaA* mutations affect central metabolic pathways

The preceding part of our metabolic analysis highlighted only one factor of our two-factorial study, namely, the change in illumination and the respective photoautotrophy to heterotrophy transitions of metabolism. However, the attenuation of the L/D response ratios compared to those of the wild type can have multiple alternative causes. Among other causes, the loss of L/D responsiveness can be explained by a deficiency to accumulate a metabolite pool that should increase upon illumination or a failure to decrease metabolites upon D transition that are high in L. The latter explanation applied in the analysis of the Δ*rpaA* mutant to a large set of metabolites from L/D-group I (**Figure 5**, relative-abundance section). Specifically, a set of amino acids, including alanine and the branched chain amino acids valine, isoleucine, or leucine, retained high concentrations upon the D transition (**Figure 5**). Similarly, 2PG lost L/D responsiveness in both the Δ*rpaA* and the Δ*kaiAB1C1* mutants, but Δ*kaiAB1C1* showed a nonsignificant illumination-independent increase compared to the extreme and highly significant accumulation of 2PG in the Δ*rpaA* mutant (**Figure 7**). In the following section, we specifically compared the mutant defects with the wild-type defects (**Supplemental Table S2**), thereby focusing on pathways that contain metabolites with large concentration changes to identify the fundamental metabolic aspects associated with the mutant defects.

**Figure 7.**
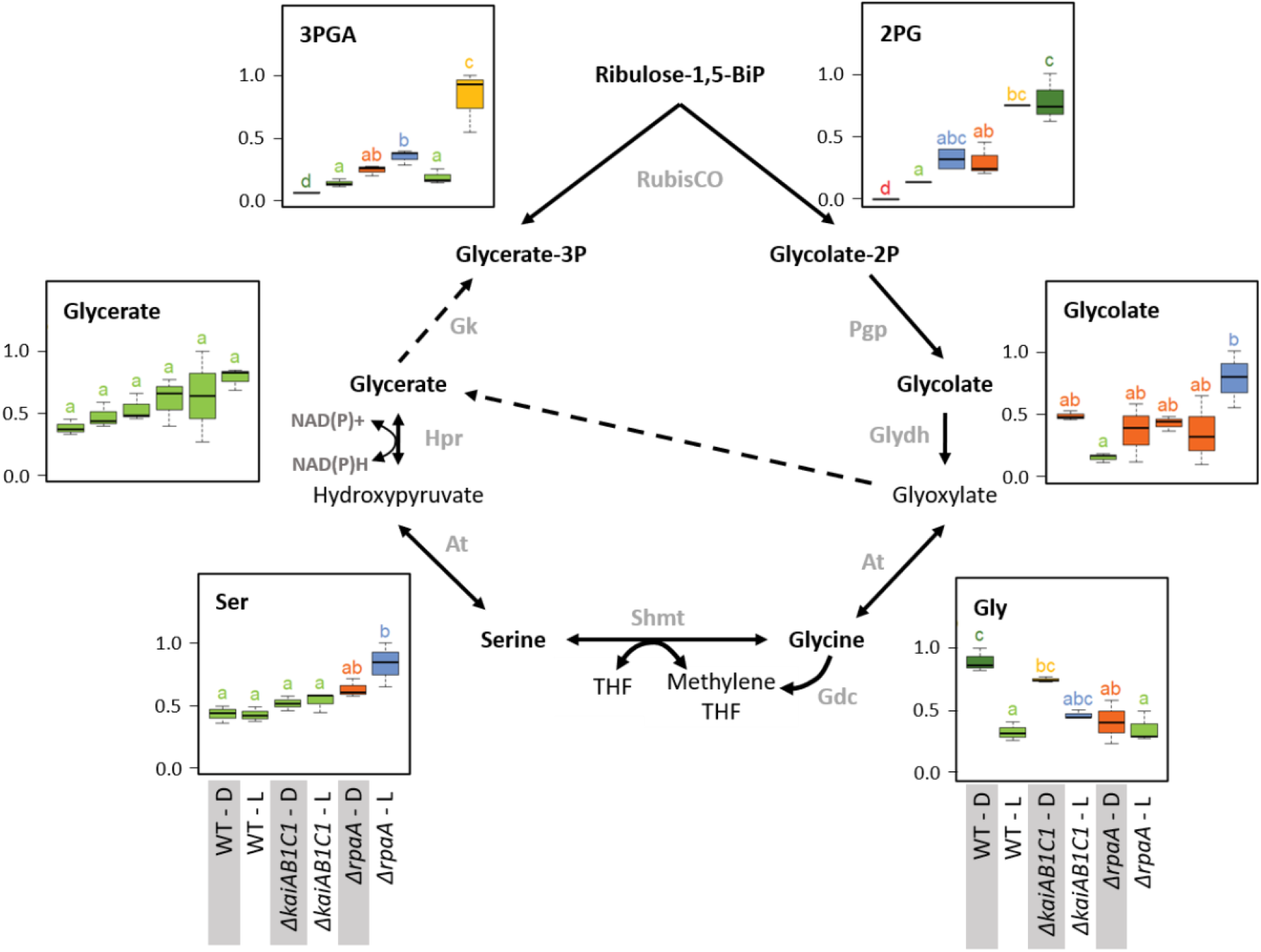
Photorespiration. Box plots showing the maximum normalized relative changes in the metabolite concentrations and Tukey test results (*P* < 0.05, n = 3). Metabolite levels that significantly differ are indicated by lowercase letters and color coding. Note the moderate and strong increases in the RubisCO products 3PGA and 2PG in the Δ*kaiAB1C1* mutant and the Δ*rpaA* mutant, respectively. High 2PG accumulation in the Δ*rpaA* mutant continues in the D phase and is associated with glycolate accumulation in the light phase. Serine and glycine respond inversely to RpaA deficiency. Glycine fails to accumulate in the D, while serine accumulates in the L. Dashed arrows indicate multiple reaction steps. Relevant enzymes and complexes include (gray) glycolate-2P phosphatase (Pgp), glycolate dehydrogenase (Glydh), aminotransferase (At), glycine decarboxylase complex (Gdc), serine hydroxymethyltransferase (Shmt), tetrahydrofolate (THF), hydroxypyruvate reductase (Hpr), and glycerate kinase (Gk).

#### Lower glycolysis

The two products of RubisCO, namely, the carboxylation product 3PGA and the oxygenation product 2PG, were highly elevated in the L samples of the Δ*rpaA* mutant and, to a lesser extent, in the Δ*kaiAB1C1* mutant (**Figure 7**). These increases suggested enhanced RubisCO activity. To support this hypothesis, we analyzed the amount of RubisCO in the mutants by a Western blot analysis (**Figure 8**). Compared to the wild type, the Δ*kaiAB1C1* and Δ*rpaA* cells accumulated more of the RbcL subunit in L and D, suggesting that the higher amounts of 3PGA and 2PG could indeed be based on a cellular increase in the RubisCO concentration. The higher amounts of 3PGA in the Δ*rpaA* mutant were consistently associated with lower glycolysis with accumulating 2PGA and PEP concentrations in this strain. These findings suggest an activation of carbon utilization from the CBB cycle in the direction of lower glycolysis and towards the oxidative branch of the TCA cycle in the absence of RpaA.

**Figure 8.**
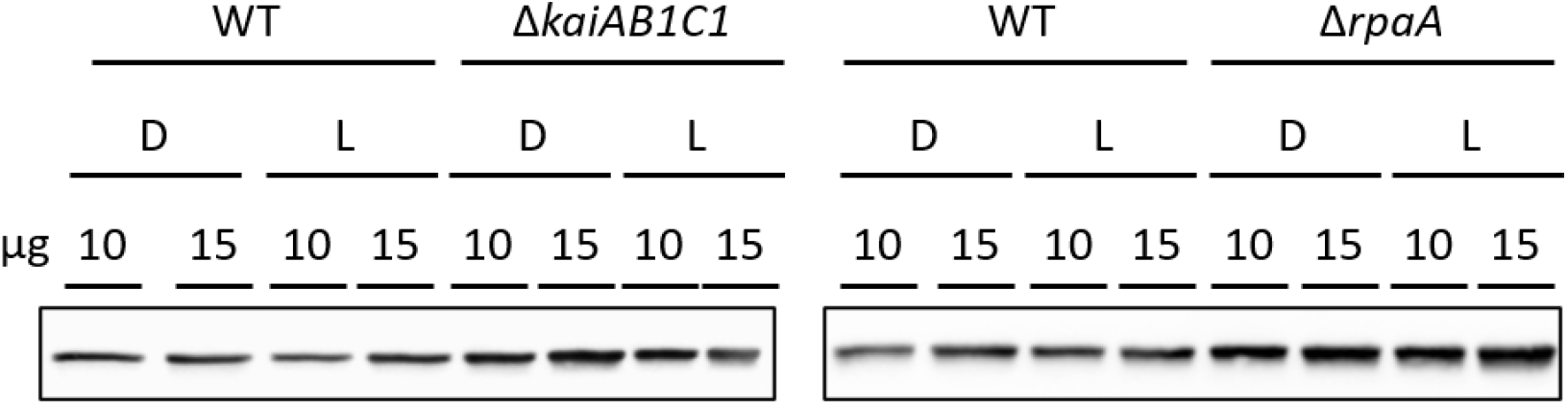
Increased RubisCO abundance in mutants compared to wild type (WT). Comparison of the RubisCO levels via immunoblot analysis using anti-RbcL. 10 or 15 μg of crude cell extract were used for SDS-PAGE. The samples were collected 5.5 h after the transition to D or L as shown in **Figure 4A**.

The assumption of the preferred use of 3PGA in the illuminated Δ*rpaA* mutant by lower glycolysis and not by gluconeogenesis was supported by the day levels of soluble carbohydrates, such as sucrose, which were not significantly changed in the Δ*rpaA* cells (**Supplemental Table S1**). Similar changes in lower glycolysis and carbohydrate metabolism occur in *Synechocystis* cells shifted from high-to low-CO_2_-supply (Orf et al., 2015). The imbalance of carbon use in favor of lower glycolysis may be controlled via the newly discovered PII-regulated switch of phosphoglycerate mutase (PGAM) activity via PirC (Orthwein et al., 2021). Previous gene expression studies implied that PirC (Sll0944) is downregulated in the Δ*rpaA* mutant under L conditions but not in D, although these data do not meet our significance criteria (Köbler et al., 2018). Hence, the lower PGAM inhibition in the Δ*rpaA* mutant likely stimulates the higher flux to 2PGA, PEP, and pyruvate compared to the wild type.

The disbalanced carbon utilization in the Δ*rpaA* cells extended to the pyruvate family of amino acids, i.e., alanine, and the branched chain amino acids leucine, isoleucine and valine. Hence, pyruvate is crucial for metabolism and cannot be depleted without affecting cyanobacteria growth (Kopka et al., 2017). According to its high demand during growth in the L, the pyruvate levels declined in the wild type upon transition from D to L. The pyruvate decrease was consistent with associated increases in the pyruvate amino acid family in the L (**Supplemental Table S1**). These amino acid responses remained unaltered in the Δ*kaiAB1C1* cells, but alanine, leucine, isoleucine and valine consistently overaccumulated in the darkened Δ*rpaA* cells (**Figure 9**). The deregulation of amino acid biosynthesis in the Δ*rpaA* mutant can contribute to the observed pyruvate depletion and may ultimately affect growth.

**Figure 9.**
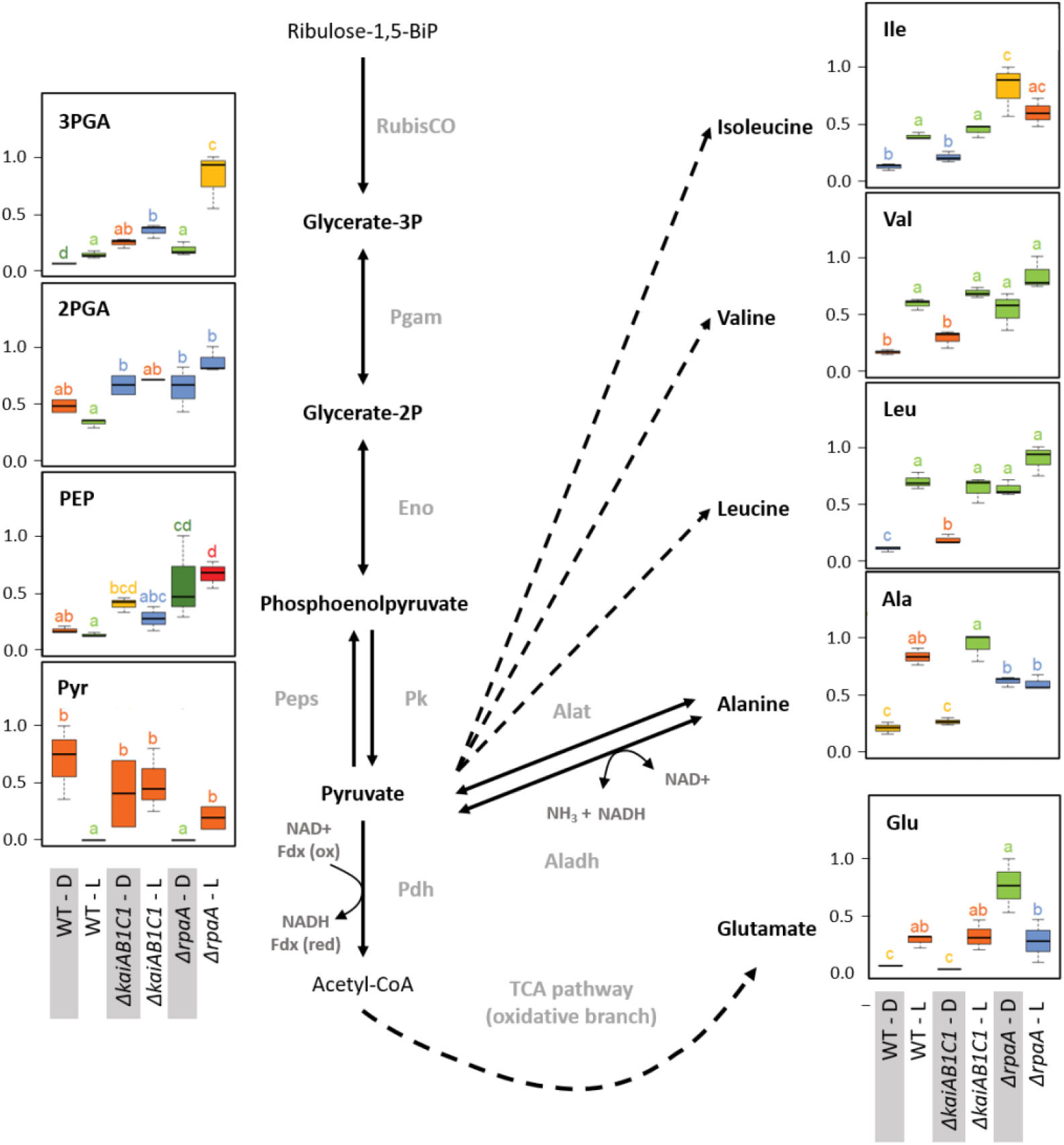
Lower glycolysis, glutamate, and the pyruvate family of amino acids. Box plots showing the maximum normalized relative changes in the metabolite concentrations and Tukey test results (*P* < 0.05, n = 3). Metabolite levels that significantly differ are indicated by lowercase letters and color coding. Note the subtle changes in the Δ*kaiAB1C1* mutant, while the Δ*rpaA* mutant exhibited significantly increased pyruvate family amino acids, 3PGA, PEP and glutamate in D. These changes are accompanied by pyruvate depletion in the D phase relative to the wild type. In the L, 3PGA accumulates in the Δ*kaiAB1C1* mutant relative to the wild type. In the Δ*rpaA* mutant, 3PGA accumulates to even higher levels, and the 2PGA and PEP increases become significant. Dashed arrows indicate multiple reaction steps. Relevant enzymes and complexes include (gray) ribulose-1,5-bisphosphate carboxylase-oxygenase (RubisCO), phosphoglycerate mutase (Pgam), enolase (Eno), phosphoenolpyruvate synthase (Peps), pyruvate kinase (Pk), pyruvate dehydrogenase (Pdh), alanine dehydrogenase (Aladh), and alanine transaminase (Alat).

#### Photorespiration

The previously mentioned increased amounts of photorespiratory 2PG extended within the pathway towards higher amounts of glycolate and serine but not glycine in the illuminated Δ*rpaA* cells (**Figure 7**). Glycine lost L/D responsiveness in the Δ*rpaA* cells, and the balance of the NADH-producing glycine decarboxylase complex (GDC) and serine hydroxymethyltransferase (SHMT) reaction favored serine. Photorespiratory metabolism was not significantly changed in the Δ*kaiAB1C1* mutant, which is consistent with the only marginal increase in 2PG in this mutant. The increased 2PG and glycolate levels indicate an increased flux into the *Synechocystis* photorespiratory pathway (Eisenhut et al., 2008; Huege et al., 2011) that detoxifies the critical substrate 2PG. The growth defect of the Δ*rpaA* cells in the L/D cycle is clearly associated with constitutively elevated 2PG levels. This observation is consistent with the finding that 2PG inhibits several key enzymes in plant primary carbon metabolism (e.g., Flügel et al., 2017). Furthermore, *Synechocystis* uses 2PG accumulation as a metabolic signal that activates the expression of cyanobacterial carbon-concentrating metabolism (CCM) because 2PG stimulates the activator CmpR and inactivates the repressor protein NdhR (Nishimura et al., 2008; Jiang et al., 2018). CCM components, such as *sbtB*, *ndhD3, ndhF3, cupA, cmpA*, and *cmpB* (Shibata et al., 2001, 2002; Xu et al., 2008; Orf et al., 2015), are consistently expressed at higher mRNA levels during the L phase in Δ*rpaA* cells (Köbler et al., 2018).

Consistent with the previously reported deregulation of photosystem I and phycobilisome gene expression in the D phase (Köbler et al., 2018), the 2PG levels largely remained at increased levels in the nonilluminated Δ*rpaA* mutant cells. In contrast, the amount of 3PGA consistently decreased in the D phase because inorganic carbon assimilation halted. RubisCO is assumed to be inactive during the night, but other sources of 2PG synthesis seem unlikely. An explanation of mistimed 2PG accumulation is proposed by the hypothesis that RubisCO may not be fully inactivated in Δ*rpaA* mutant cells during the night, especially because there seems to be substantial overaccumulation of this enzyme in these cells (**Figure 8**). RubisCO which is not completely inactivated may then preferentially perform oxygenation during the night due to the low C_i_/high O_2_ intracellular environment. This explanation not only applies to the mutant cells but also seems to apply to the wild type, which fully detoxifies 2PG in D but still contains significantly higher amounts of glycolate and glycine at night compared to the day (**Figure 7**). An alternative explanation of elevated 2PG in D may be a potential carry-over of the high concentration from the preceding L phase. Such an assumption could imply D inhibition of the 2PG dephosphorylation step in the Δ*rpaA* mutant. However, regulation of the 2PG phosphatase step seems unlikely because of the presence of up to four partially highly promiscuous 2-PG phosphatases within the *Synechocystis* genome (Rai et al., 2018).

#### TCA pathways

Our metabolome analysis covered most metabolites of the noncanonical TCA cycle (**Figure 10**) and associated amino acid biosynthesis pathways that receive carbon building blocks from this cycle (**Figure 11**). Among cyanobacteria, the TCA cycle is not closed because the 2-oxoglutarate (2OG) dehydrogenase complex is missing. The open nature of the TCA pathways is clearly reflected in our current study by the inverted L/D responses of the TCA cycle metabolites from its reductive (C4) and oxidative (C6 and C5) branches. Malate, fumarate and succinate of the reductive branch decreased in L with large decreases in the malate and fumarate pools in all strains but only minor changes in succinate. This consistent L/D response indicates carbon utilization by respiration via the succinate dehydrogenase complex in D and the inactivation or reversion of this process in L. In contrast, the intermediates of the oxidative branch, such as citrate and aconitate, accumulated in L and reflected enhanced use of photosynthetically fixed organic carbon by ammonia assimilation via glutamine synthetase/glutamine:2-oxoglutarate aminotransferase (GS/GOGAT cycle). However, the amount of 2OG did not significantly differ between the L and D samples of all strains, ruling out the possibility that 2OG-mediated signaling was responsible for the observed expression changes in the CCM-related genes. The deficiency of the KaiAB1C1 system and the RpaA regulator did not affect the TCA pathways, but the utilization of carbon in the associated nitrogen assimilation is reported in the following section.

**Figure 10.**
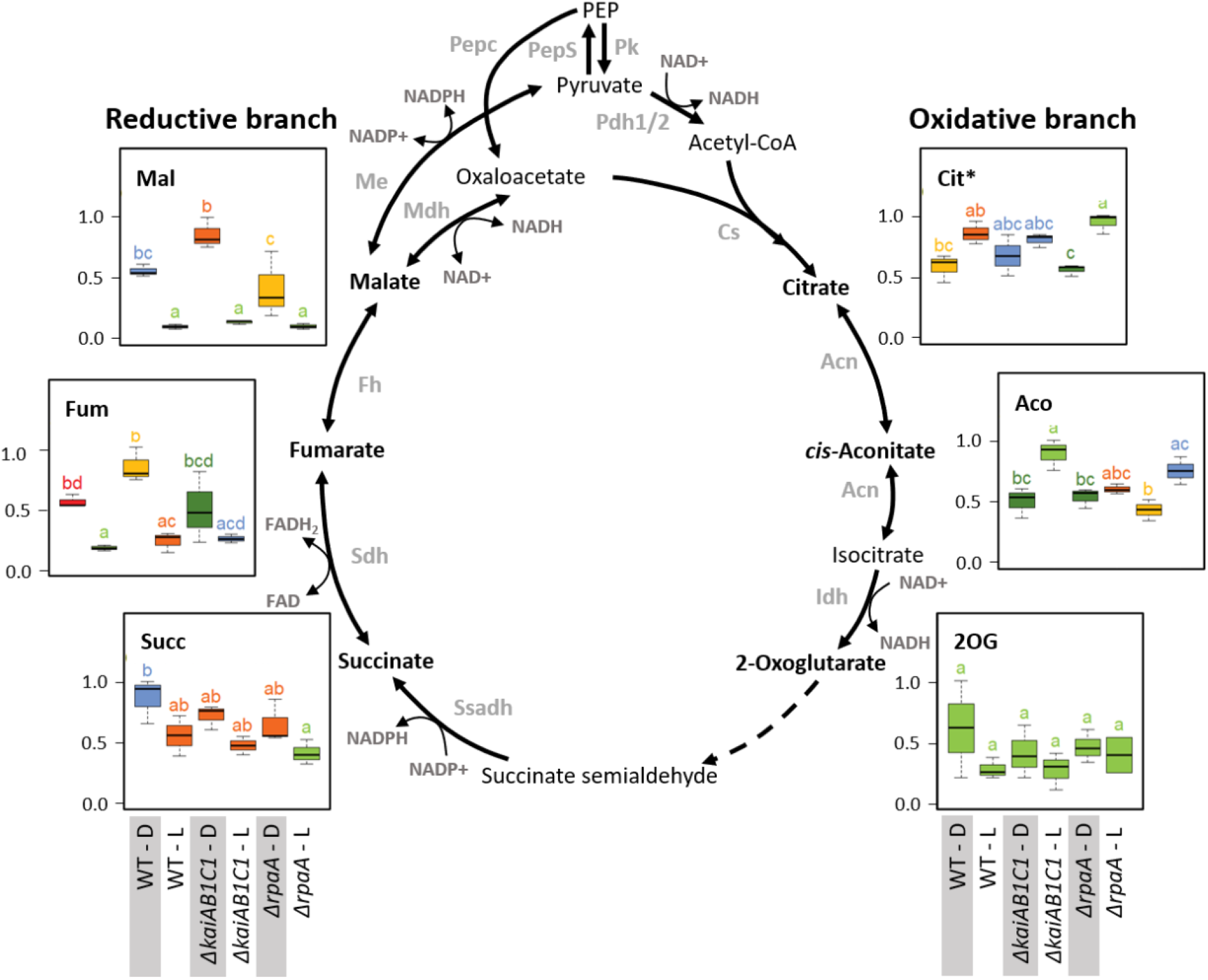
Tricarboxylic acid pathways. Box plots showing the maximum normalized relative changes in the metabolite concentrations and Tukey test results (*P* < 0.05, n = 3). Metabolite levels that significantly differ are indicated by lowercase letters and color coding. Note the inverse accumulation of metabolites from the oxidative and reductive branches of the TCA pathways. The concentrations of the intermediates of the TCA reactions were largely unchanged in the Δ*kaiAB1C1* and Δ*rpaA* cells. Dashed arrows indicate multiple reaction steps. Relevant enzymes and complexes included (gray) phosphoenolpyruvate synthase (PepS), pyruvate kinase (Pk), pyruvate dehydrogenase (Pdh), phosphoenolpyruvate carboxylase (Pepc), citrate synthase (Cs), aconitase (Acn), isocitrate dehydrogenase (Idh), succinate semialdehyde dehydrogenase (Ssadh), succinate dehydrogenase (Sdh), fumarase (Fh), malate dehydrogenase (Mdh), and malic enzyme (Me).

**Figure 11.**
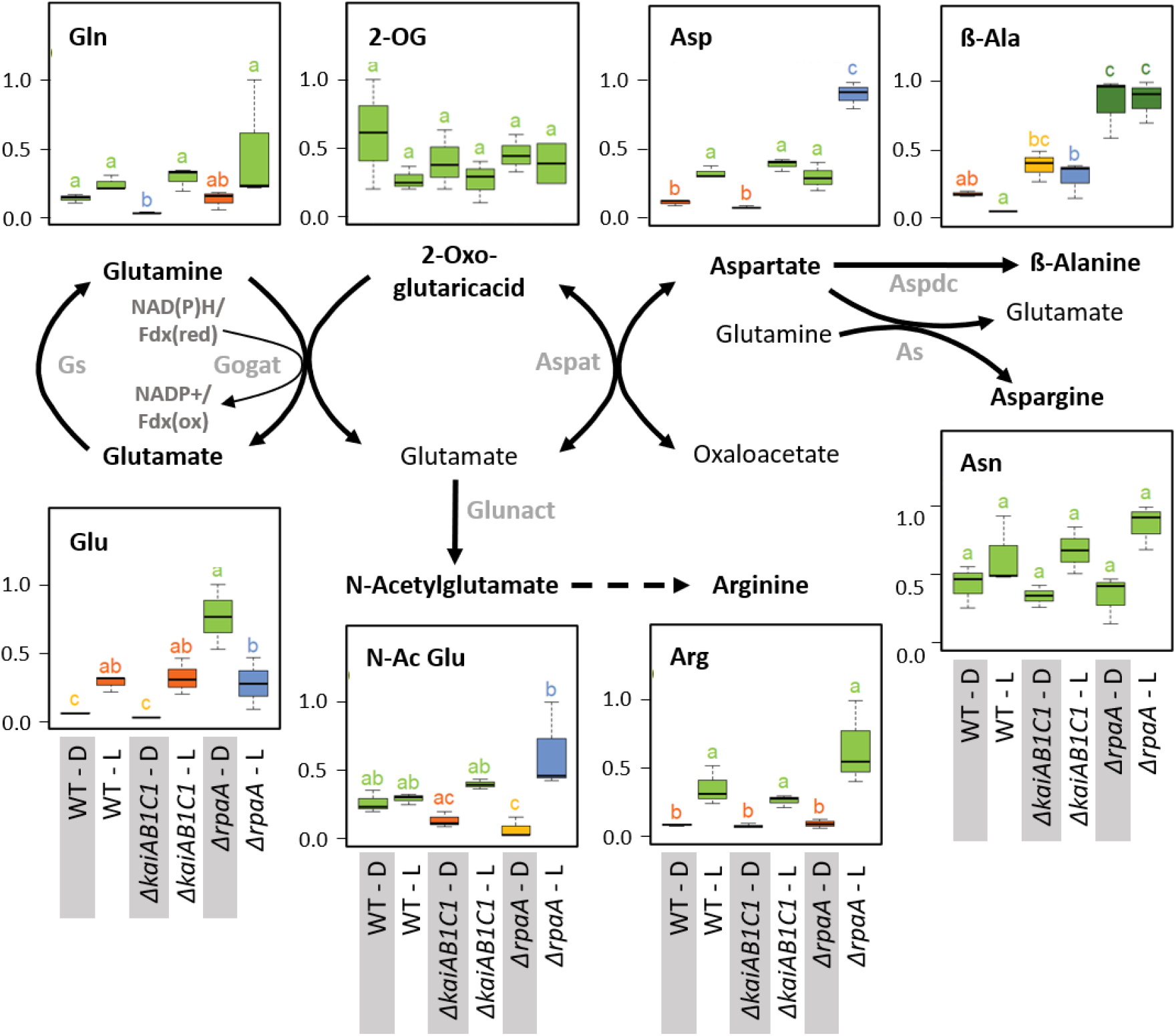
Nitrogen assimilation and associated amino acid biosynthesis. Box plots showing the maximum normalized relative changes in the metabolite concentrations and Tukey test results (*P* < 0.05, n = 3). Metabolite levels that significantly differ are indicated by lowercase letters and color coding. Note the increase in aspartate and the D accumulation of glutamate (**Figure 11**) in the Δ*rpaA* mutant. Relevant enzymes and complexes include (gray) glutamine synthetase (Gs), glutamine 2-oxoglutarate aminotransferase (Gogat), aspartate aminotransferase (Aspat), asparagine synthetase (As), aspartate decarboxylase (Aspdc), and glutamate N-acetyltransferase (Glunact).

#### Nitrogen assimilation and amino acid biosynthesis

In addition to 2OG, the products of the GS/GOGAT cycle, i.e., glutamine and glutamate, and the subsequent aminotransferase product aspartate accumulate in illuminated wild-type cells (**Figure 11**). The accumulation of these amino acids paralleled the changes in the oxidative TCA branch. These concentration changes are consistent with the operation of ammonia assimilation in the L. Nitrogen (N)-assimilation requires the products of photosynthetic light reactions, such as ATP and reduced ferredoxin. Similar to the TCA pathways, the deficiency of KaiAB1C1 proteins did not affect ammonium assimilation or the associated glutamate/aspartate metabolism. The Δ*rpaA* mutant, however, differed in the L/D responses of glutamate but not of glutamine; specifically, the glutamate concentrations increased in D (**Figure 11**). This increase was associated with a significant accumulation of aspartate in the D and consistent with the supply directly from glutamate via the aspartate aminotransferase reaction, while the carbon backbone, oxaloacetate, may be supplied by phosphoenolpyruvate carboxylase (PEPC), malate dehydrogenase and the intermediates of the reductive TCA branch. The lack of a concomitant accumulation of 2OG indicated active reuse by the GS/GOGAT cycle. Upon illumination, the Δ*rpaA* cells overaccumulated Asp and N-acetylglutamate, which is an additional direct product of glutamate in the pathway towards arginine biosynthesis (**Figure 11**). Collectively, our observations indicate overly active N assimilation in Δ*rpaA* cells that extends in D to multiple proteinogenic amino acid pools (**Figure 5**). Furthermore, in both diurnal phases, the amounts of nonproteinogenic amino acids, such as β-alanine and homoserine, increased, while nucleobases, such as thymine and uracil, increased in L (**Figure 11**, **Supplemental Table S2**). We hypothesize that the Δ*rpaA* mutant may not stop N-assimilation during the night for an unknown reason.

#### Metabolic cofactors

The set of nucleotides and nucleosides monitored in the current study mostly exhibited minor pool changes that did not pass the significance threshold (**Supplemental Table S1, Supplemental Table S2**). The lack of significance in our study likely results from the high variation in the sampling time in the middle of the L phase. This phase still represents a time during which the concentrations of many nucleotides and nucleosides return from high levels in the morning (Saha et al., 2016) to the low levels present at the end of the day and during the night (Werner et al., 2019).

The levels of glutathione (GSH) and flavin mononucleotide (FMN) fluctuated less. Both metabolites indicate the deregulation of the redox state in Δ*kaiAB1C1* cells and greater deregulation in the Δ*rpaA* mutant. GSH failed to increase upon illumination in the Δ*rpaA* cells, whereas FMN accumulated constitutively in both the Δ*kaiAB1C1* and Δ*rpaA* cells.

Motivated by these observations, we monitored the nicotinamide adenine dinucleotide redox cofactors NADP^+^, NADPH, NAD^+^ and NADH. NADP^+^, NADPH and NAD^+^ were quantified, but these nucleotides and nucleosides did not pass our significance threshold (**Supplemental Table S3**). NADH was not detectable due to low concentrations. Our measurements indicated that NADP^+^, NADPH, NAD^+^, and the NADPH/NADP ratio potentially increased in the wild type in the L phase compared to D. This increase in the NADPH/NADP ratio in L appeared to be absent from the Δ*kaiAB1C1* and Δ*rpaA* mutant cells. In both mutants, this observation can be explained by an increase in the NADPH levels in the D (**Supplemental Table S3**).

## 4 Discussion

Based on metabolome analyses of *Synechocystis* mutants in L and D samples of autotrophically grown cultures, we can conclude that the *Synechocystis* KaiAB1C1-SasA-RpaA potential timing system largely affects the metabolic switch from photoautotrophy during the day to heterotrophy at night. This effect is comparable to the function of the *Synechococcus* sp. PCC 7942 circadian clock system, although the effects of the clock deficiency on the transcriptomic and metabolomic levels differ between these two cyanobacteria. These strain-specific differences are likely caused by differential adaptation of the two cyanobacteria. Facultative heterotrophy and the presence of alternative putative clock components in *Synechocystis* inevitably impact the transcriptional regulatory network with divergent metabolic consequences in the two cyanobacterial strains. Our findings suggest that the molecular mechanisms unraveled in one model organism may differ from those in other strains due to adaptive evolution. This assumption is consistent with findings suggesting that paralog duplications may quickly vanish from cyanobacterial genomes, e.g., of *Acaryochloris* or *Chlorogloeopsis* strains, unless they provide fitness benefits (Miller et al., 2011; Weissenbach et al., 2017). This reasoning likely applies to clock oscillator-related genes; for instance, the *Nostoc linckia kaiABC* gene family has been shown to respond to long-term differential stress by rapid adaptive radiation (Dvornyk et al., 2002).

In our *Synechocystis* model, the *rpaA* deletion had a much larger effect on metabolism than the *kaiAB1C1* mutation, which is consistent with the more pronounced transcriptional changes in Δ*rpaA* (Köbler et al., 2018), although both mutants grew very similarly under these conditions. Most interestingly, Δ*rpaA* showed a remarkable overaccumulation of 2PG and amino acids in D with concurrent effects on the redox balance. The effects in the Δ*kaiAB1C1* mutant are similar but smaller. This can be mainly explained by RpaA function in transcriptional regulation even in the absence of Kai-controlled phosphorylation. Phenotypic analyses of phosphor-mimicking variants of RpaA could be helpful to evaluate this hypothesis in future studies. Further, the lack of RpaA-RpaB interactions in the Δ*rpaA* mutant might affect the control of the large RpaB-dependent regulon, which partially overlaps with the RpaA regulon (e.g., control of the *psaAB* operon) (Seino et al., 2009; Köbler et al., 2018). Moreover, a recent study by Riediger et al. (2019) largely extended the RpaB regulon, which currently includes many enzymes involved in primary carbon and nitrogen metabolism, likely contributing to the larger impact of RpaA deficiency on L/D-mediated metabolic changes. In addition, or alternatively, the presence of the likely only partially neofunctionalized KaiB3C3 and KaiB2C2 systems in *Synechocystis* could compensate for the loss of the KaiAB1C1 complex. At least for the KaiC3-based system, it has been shown that it interacts with the KaiAB1C1 complex and modulates its function (Wiegard et al., 2020).

It has been suggested that the NADPH/NADP ratio plays a major role in the diurnal growth of *Synechocystis* (Saha et al., 2016). This ratio is known as the main redox signal and has a pronounced impact on enzyme activities in the CBB cycle, glycolysis and gluconeogenesis. For example, the NADPH/NADP ratio switches the activities of the CBB enzymes Gap2 and Prk on and off via the small redox-sensitive protein CP12 in L to D transitions (McFarlane et al., 2019). Hence, the tendency of higher NADPH/NADP ratios in the mutants, especially in Δ*rpaA*, suggests that these and other metabolic redox cofactors likely contribute to the observed disbalanced carbon and nitrogen metabolism in L/D transitions. Moreover, the continued and enhanced activity of RubisCO in D leads to the observed accumulation of 2PG in both mutants due to the enhanced oxygenase reaction in the presence of lower C_i_/O_2_ ratios in D. 2PG can be toxic and provides a metabolic signal that falsely indicates carbon limitation. 2PG accumulation in Δ*rpaA* is much more pronounced than that in Δ*kaiAB1C1*, explaining the previously detected overexpression of transcripts involved in C_i_ limitation response in Δ*rpaA* (Köbler et al., 2018). The PHB deficiency observed in both strains can be traced to the lower accumulation of mRNAs encoding the enzymes involved in PHB synthesis. Furthermore, this deficiency might be caused by the lower pyruvate pool and respective metabolic limitation of the acetyl-CoA precursor of PHB synthesis in the clock mutants. However, the lack of PHB should not affect the growth of cells in L/D cycles as shown by Damrow et al. (2016), suggesting that this deficiency is not the primary cause of the phenotype of the analyzed mutants.

Moreover, previous analyses showed that mixotrophic conditions, i.e., feeding glucose in the L, have an aggravating effect on the growth of both mutant strains (Dörrich et al., 2014; Köbler et al., 2018). In the present study, we provide evidence suggesting that high CO_2_ supplementation improves the ability of both mutants to grow at diurnal L/D cycles. In contrast, glucose addition and limiting C_i_ availability result in redox imbalance. Therefore, the defect in the mutants’ ability to grow diurnally is likely associated with disturbed redox homeostasis as previously shown in *Synechococcus elongatus* PCC 7942 (Diamond et al., 2017). The redox imbalance in the two mutants is also reflected by the transcriptional changes as genes encoding flavoproteins (Flv), especially Flv2 and Flv4, are significantly upregulated compared to those in the wild type (Köbler et al., 2018). These flavoproteins along with Flv1 and Flv3 are required for the Mehler-like reaction in *Synechocystis* to support growth under fluctuating or high light conditions (Santana-Sanchez et al., 2019).

Finally, motivated by the potential of *Synechocystis* and other cyanobacteria for biotechnological applications and the important role of the circadian clock in controlling cyanobacterial metabolism, several groups utilized deletion and overproduction strains of the SasA-RpaA output module and resulting analyzed metabolic changes. Most importantly, the overexpression of *sasA* and *rpaA* resulted in enhanced sugar catabolism in the D (Iijima et al., 2015; Osanai et al., 2015). Our analysis of a mutant lacking the *rpaA* gene reveals the opposite effects, i.e., more carbon is exported from the CBB cycle into lower glycolysis and TCA cycle, which is consistent with published data of *rpaA* overexpression (Iijima et al., 2015). Hence, the results presented here and by other groups provide evidence suggesting that an intact KaiAB1C1-based system is required in *Synechocystis* for the improved production of C-based metabolites under natural L/D cycles with cyanobacteria.

## 5 Author Contribution

A.W., J.K., and C.K. designed research; N.M.S., Y.R., S.T, and C.K. performed research; N.M.S., Y.R., J.K., and M.H. analyzed the data; and J.K., M.H., A.W., N.M.S., and Y.R. wrote the paper.

## 7 Acknowledgment

We thank Matthias Boll for providing the UPLC and Max Willistein for support in the operation of the UPLC. We further thank Judith Asal and Werner Bigott for excellent technical assistance.

## Footnotes

1 http://chemdata.nist.gov/

2 https://rstudio.com/

