## Supplementary figures for "Homologs of circadian clock proteins impact the metabolic switch between light and dark growth in the cyanobacterium *Synechocystis* sp. PCC 6803"

### *Supplementary Material*

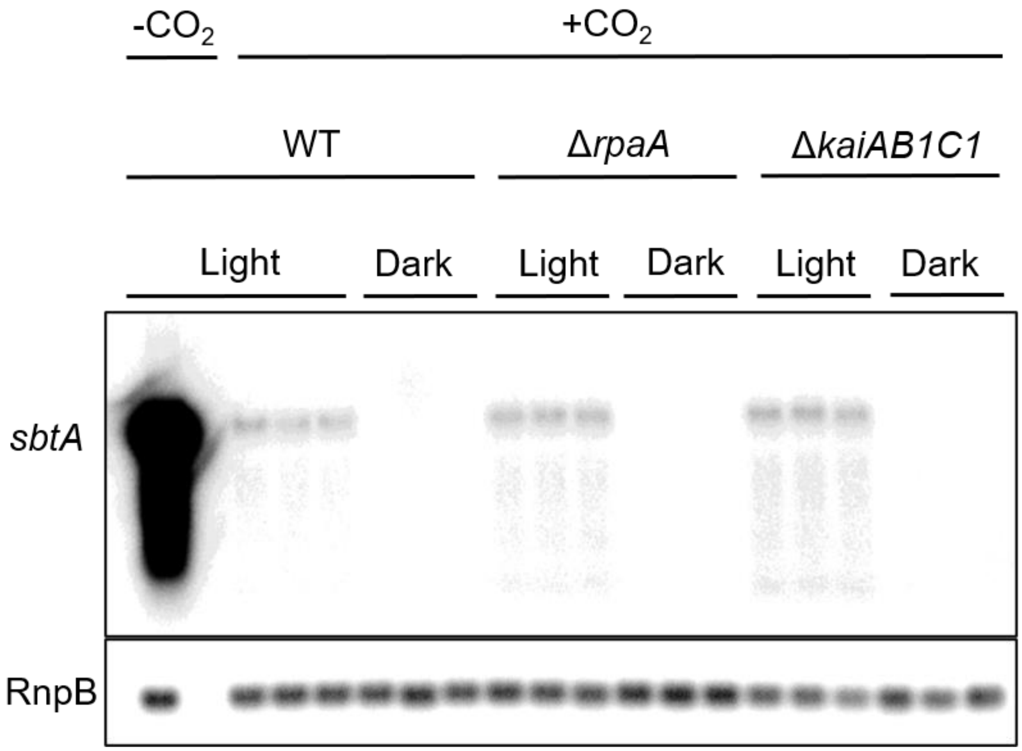


**Supplementary Figure 1. *ΔrpaA* and *ΔkaiAB1C1* strains show no $C_i$ response at high $CO_2$ concentrations.** Northern blot analysis of *sbtA* mRNA accumulation in wild-type (WT) and clock mutant strains under L/D conditions with 1% $CO_2$. RNA from three independent cultures was isolated in the middle of the L and D phases. Hybridization was performed with a *sbtA*-specific RNA probe, and RnpB hybridization was used as a loading control. RNA of wild type cultivated under ambient air was used for comparison.

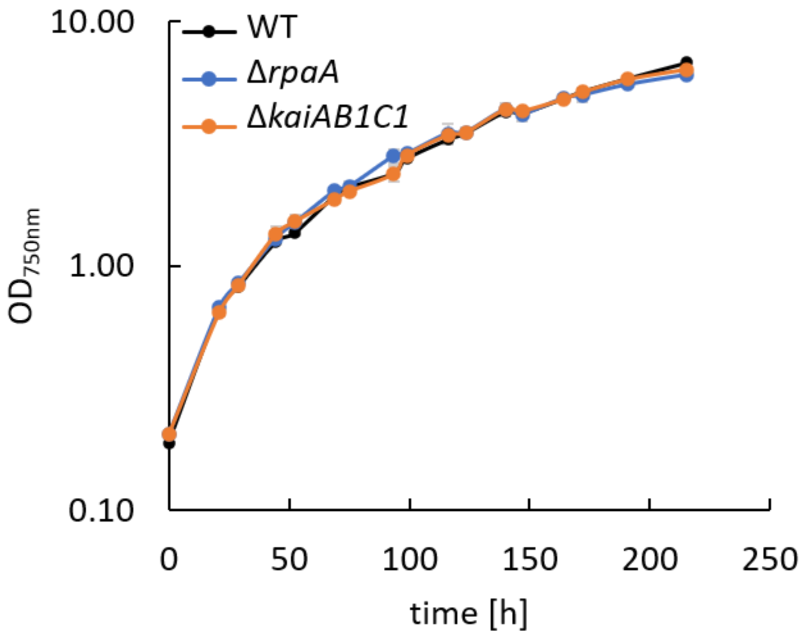


**Supplementary Figure 2. Growth of *Synechocystis* clock mutants under constant light conditions.** Wild-type (WT) and Δr*paA* and Δ*kaiAB1C1* deletion mutants were grown under continuous illumination with 75 µmol photons m$^{-2}$s$^{-1}$ in ambient air. Each point represents the mean of three technical replicates (± standard deviation).

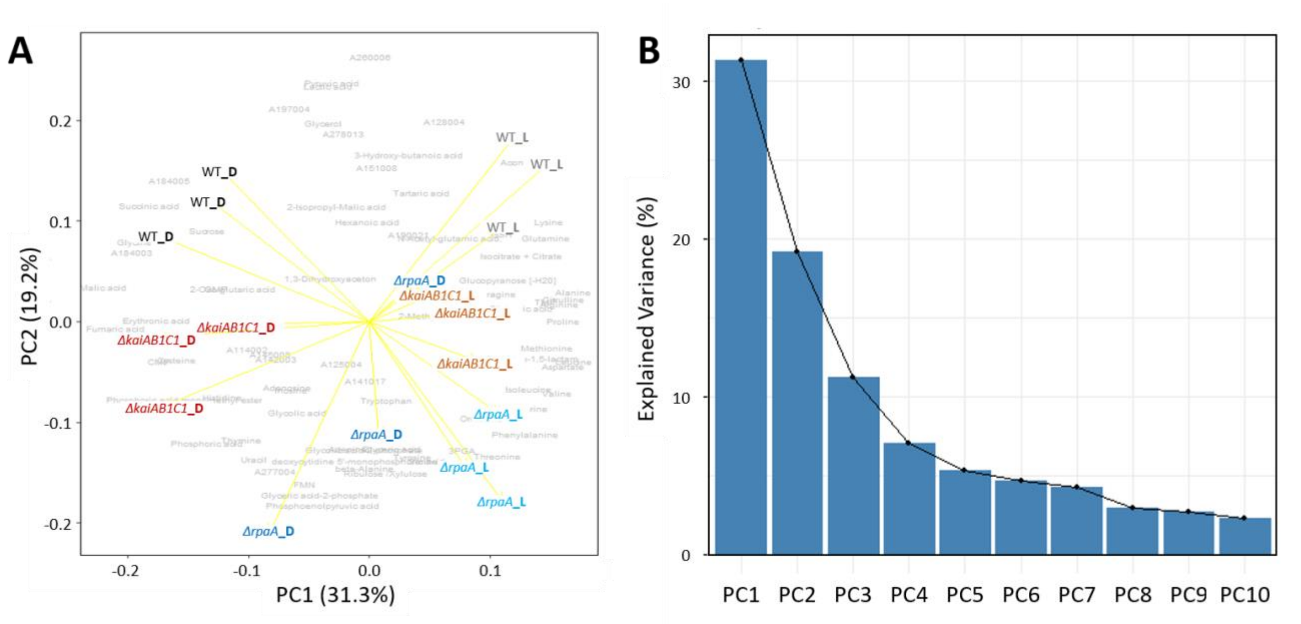


**Supplementary Figure 3. Principal component analysis (PCA) of the metabolic light dark switch in the *ΔkaiAB1C1* clock oscillator and the *ΔrpaA* output kinase mutants compared to the *Synechocystis* wild type.** (A) Biplot of PC1 and PC2. (B) Scree plot of the variance contribution to PC1-PC10. This supplement corresponds to Figure 4, and the details are provided in its legend.

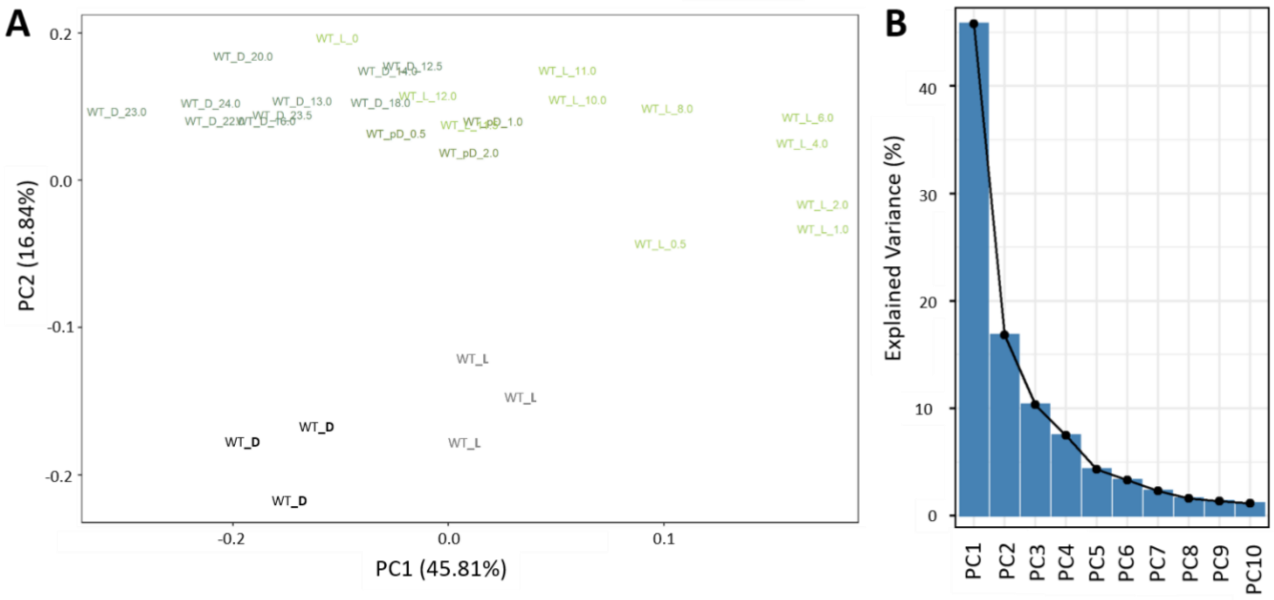


**Supplementary Figure 4. Principal component analysis (PCA) comparing the metabolic light and dark states of the *Synechocystis* wild type between this study and a previous diurnal analysis** (Werner et al., 2019). PCA combines 47 common metabolites from the two studies that were separately maximum scaled prior to the data merging. The merged data matrix was log-transformed and autoscaled prior to PCA. PC1 indicates high agreement of the light/dark changes between the studies. PC2 describes the differences caused by deviating cultivation conditions. Note that an alignment to specific diurnal phases between the studies was impossible.

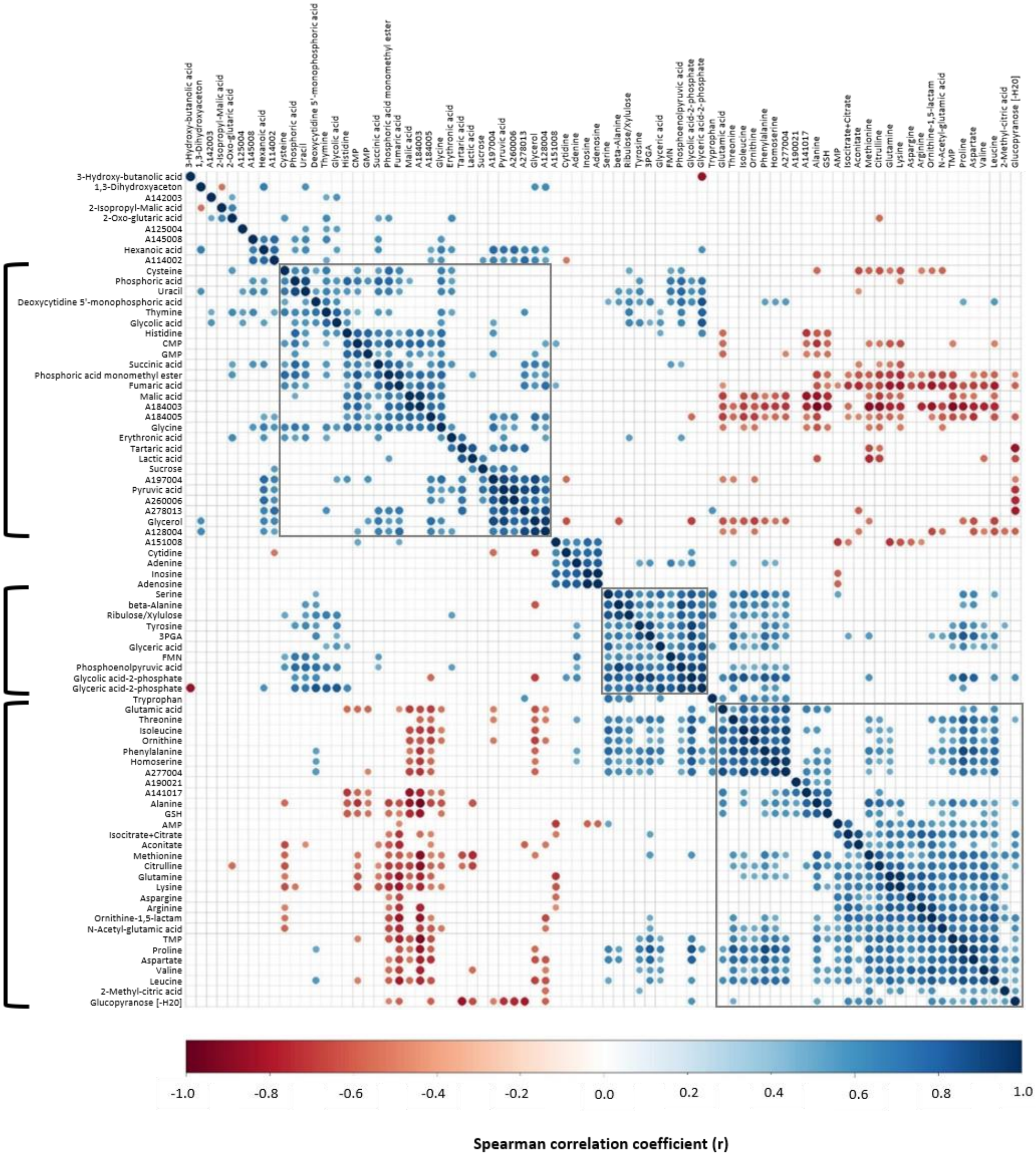


**Supplementary Figure 5. Global correlation matrix of all metabolites monitored in this study.** All annotated LC- and GC-MS metabolite data were combined after separate maximum scaling. Spearman's correlations among metabolites were calculated using all mutant and wild-type data, including both illumination conditions. The resulting symmetric matrix was arranged by hierarchical clustering using the Euclidian distance to yield groups of highly intercorrelated metabolites across the diagonal. The plot shows only significant correlations (*P* < 0.05). The color scale indicates Spearman´s correlation coefficient (r) in the range of +1.0 (blue) to -1.0 (red), noncorrelated (white). The brackets to the left indicate a small central group of predominantly positively correlated metabolites and two groups (top and bottom) of metabolites that are predominantly positively correlated within the group and partially negatively correlated between the two groups.
